# Species-level controls of foliar methane and nitrous oxide fluxes: roles of traits and microbes in temperate trees

**DOI:** 10.64898/2026.04.06.716737

**Authors:** Md Rezaul Karim, Sean C. Thomas

## Abstract

The contribution of tree foliage to atmospheric methane (CH_4_) and nitrous oxide (N_2_O) fluxes remains a major uncertainty in global GHG budgets. We made repeated in situ measurements of foliar CH_4_ and N_2_O fluxes across 25 temperate tree species interplanted at a forest restoration site using high-resolution laser spectroscopy. Tree foliage was consistently a net CH_4_ sink and a net N_2_O source in all species. Foliar CH_4_ oxidation increased by ∼33% in fall relative to spring and was ∼3-fold higher in shade-tolerant than shade-intolerant angiosperm species. Species differences accounted for most of the variability in fluxes, while correlations with soil emissions were comparatively weak. Microbial DNA sequencing revealed that the highest CH_4_-oxidizing angiosperm species (*Tilia americana*) harbored abundant Type I methanotrophs, whereas the lowest-oxidizing species (*Prunus virginiana*) had nearly 100-fold lower methanotroph abundance, with a foliar microbial community dominated by facultative methylotrophs. Global warming potential (GWP) scaling indicates that foliar CH_4_ uptake overwhelmingly dominates the net climate forcing effect. Our results suggest that the large and predictable differences in foliar CH_4_ uptake among tree species and associated differences in foliar microbial communities are of importance in understanding and potentially enhancing the global terrestrial CH_4_ sink.

## INTRODUCTION

Methane (CH_4_) and nitrous oxide (N_2_O) are highly potent biogenic greenhouse gases (GHGs) with global warming potentials 28–34 and 265–298 times greater than CO_2_, respectively, over a 100-year timescale (Pachauri *et al*., 2014; Nisbet *et al*., 2021). Despite their lower atmospheric concentrations, their high radiative efficiency and, for N_2_O, a prolonged atmospheric lifetime (∼116 years) amplify their impact on climate forcing (Prather *et al*., 2015). Since the preindustrial period, atmospheric concentrations of CH_4_ and N_2_O have increased by ∼156% and 23%, respectively, resulting in estimated radiative forcings of 0.54 W m^−2^ (CH_4_) and 0.21 W m^−2^ (N_2_O) as reported in the IPCC Sixth Assessment Report 2021 (Masson-Delmotte *et al*., 2021). While anthropogenic sources such as agriculture and fossil fuel combustion dominate global budgets, terrestrial ecosystems–including soils and vegetation–represent significant regulators of CH_4_ and N_2_O fluxes. Mechanisms of aboveground exchange, particularly via foliage, remain poorly constrained, limiting their representation in climate models.

Historically, CH_4_ and N_2_O research has focused on soil and wetland systems, where anaerobic and redox-sensitive microbial processes mediate most fluxes (Aurangojeb *et al*., 2017; Pangala *et al*., 2017; Covey & Megonigal, 2019). However, a growing body of evidence now indicates that trees, including stems and leaves, can act as both sources and sinks of CH_4_ and N_2_O (Barba *et al*., 2019; Gorgolewski *et al*., 2022, 2023; Karim *et al*., 2024). Although per-leaf-area fluxes are low, the global foliage surface area exceeds 1.5 × 10^8^ km^2^ (Asner *et al*., 2003), suggesting even modest fluxes may have substantial ecosystem-level impacts (Gauci *et al*., 2024).

Early interest in foliar CH_4_ exchange arose from a report of aerobic CH_4_ emissions from plants by (Keppler *et al*., 2006); however, this result was challenged due to methodological artefacts, with studies suggesting that UV-induced photochemical reactions contribute to only very small CH_4_ fluxes in-situ (Kirschbaum *et al*., 2006; Dueck *et al*., 2007). More recent investigations suggest that xylem-mediated transport of soil-derived CH_4_ and internal microbial oxidation are the main mechanisms determining foliar CH_4_ exchange (Putkinen *et al*., 2021; Gorgolewski *et al*., 2023; Karim *et al*., 2025). In parallel, phyllosphere microbial communities have been shown to include methanotrophic bacteria, particularly Type II taxa such as *Methylocystis* and *Methylosinus*, which colonize leaves and are capable of oxidizing CH_4_ under well-lit, oxic conditions (Iguchi *et al*., 2012, 2013). These phyllosphere methanotrophs exhibit adaptations to UV stress, desiccation, and nutrient limitation, suggesting their persistence under field-relevant conditions (Legein *et al*., 2020; Mohanty *et al*., 2023). However, while marker gene detection (e.g., pmoA) supports their presence, direct evidence of active CH_4_ oxidation remains scarce, and the functional role of phylloplane microbes vs. endophytes is still unresolved.

Similarly, until recently foliar N_2_O exchange has been overlooked but is now recognized as a potentially important flux pathway. Emissions typically dominate over uptake (Karim *et al*., 2024, 2025), and several mechanisms have been proposed. Translocation of soil-derived N_2_O via xylem flow has been demonstrated (Welch *et al*., 2019), but there is also compelling evidence for endogenous N_2_O production through internal plant processes. One pathway involves mitochondrial reduction of nitric oxide (NO) to N_2_O under hypoxic stress, serving as a detoxification mechanism (Timilsina *et al*., 2020). Recent surface sterilization experiments have shown no significant difference in foliar N_2_O emission between sterilized and non-sterilized leaves (Qin *et al*., 2024), suggesting that microbial surface processes are not an important source. Nonetheless, under natural conditions, where microbial diversity is greater and nutrient inputs vary, microbial nitrifiers or denitrifiers may still contribute. Physiological parameters such as foliar nitrogen content, stomatal conductance, and transpiration rates are likely to regulate these pathways (Karim *et al*., 2025) but remain poorly quantified across species.

Despite increasing recognition of foliar contributions to forest GHG budgets, in-situ survey studies remain limited in number, geographic scope, and taxonomic breadth (Gorgolewski *et al*., 2023; Karim *et al*., 2025). Existing studies have largely focused on tropical or wetland systems (Pangala *et al*., 2017; Karim *et al*., 2024), with temperate forests–particularly broadleaf and needleleaf tree species–receiving minimal attention. Available data are restricted to a small number of species, leaving significant gaps in our understanding of interspecific and functional trait-based variability in foliar fluxes.

Methodological limitations have also constrained progress. Early studies frequently employed static chamber plus gas chromatography techniques, which are inadequate for resolving low-level foliar gas fluxes and prone to artifacts under fluctuating environmental conditions (Takahashi *et al*., 2012). Recent advances in dynamic chamber systems coupled with laser-based spectroscopy (e.g., CRDS, OA-ICOS, TILDAS) now permit rapid sub-ppb measurements of CH_4_ and N_2_O under field conditions, enabling continuous, high-resolution flux estimates (Barba *et al*., 2019; Gorgolewski *et al*., 2023; Karim *et al*., 2024). Additionally, these systems allow analyses of temporal variation in foliar CH_4_ and N_2_O exchange–linked to diel cycles, phenological stage, and environmental conditions such as light and water availability (Kohl *et al*., 2023; Tenhovirta *et al*., 2024; Karim *et al*., 2025).

To date, microbial studies related to foliar CH_4_ exchange have relied on marker genes (e.g., pmoA) without assessing transcription or localization (Moisan *et al*., 2025), leaving endophytic contributions inferred rather than demonstrated. Existing metagenomic and marker-gene surveys are rarely integrated with field-based CH_4_ flux measurements, leaving the functional role of foliar methanotrophs unresolved. The potential role of methanotrophic endophytes provides a framework to predict seasonal and interspecific variability in CH_4_ oxidation. If colonization or proliferation progresses through the growing season, CH_4_ oxidation rates should increase from spring to autumn as endophytic communities mature. This pattern is likely to be most pronounced in deciduous species, with annually developing leaves, and less so in evergreen conifers, where needles persist for multiple years and host stable microbial assemblages (Baldrian, 2017). If methanotroph abundance drives oxidation capacity, as observed in soils (Lima *et al*., 2014) and bark (Jeffrey *et al*., 2021), species with higher foliar CH_4_ uptake should exhibit greater representation of methanotrophic taxa.

In addition to microbial influences, functional traits related to shade tolerance and successional status may predict interspecific variation in foliar CH_4_ and N_2_O exchange. Shade-tolerant, late-successional species maintain longer leaf lifespans (Seiwa *et al*., 2006), facilitating progressive colonization by methanotrophic endophytes. Although leaf lifespan of temperate-zone trees is limited by growing season length, temperate pioneer species undergo multiple leaf flushes and retain foliage for shorter periods (Kikuzawa, 2003), potentially limiting microbial establishment. Lower water fluxes in shade-tolerant species (Zhang *et al*., 2016) may also reduce xylem transport of soil-derived CH_4_, enhancing the relative role of in situ oxidation. Higher foliar nitrogen in pioneer species may increase endogenous N_2_O production, explaining lower CH_4_ oxidation and higher N_2_O emissions, consistent with tropical forest observations (Karim *et al*., 2024). These considerations guided the present study, in which we surveyed diverse temperate trees to identify variability in foliar CH_4_ oxidation and examine links with tree shade-tolerance, seasonality, and microbial composition.

Here we test the overarching hypothesis that interspecific and seasonal variation in foliar GHG exchange reflects the combined influence of plant functional traits, environmental conditions, and microbial associations. Specifically, we hypothesize that (i) foliar CH_4_ oxidation will vary predictably with species shade tolerance, being higher in shade-tolerant and lower in shade-intolerant pioneer species; (ii) within species, CH_4_ oxidation will increase from spring to autumn as methanotrophic endophytes proliferate within maturing foliage; and (iii) species exhibiting greater foliar CH_4_ oxidation will host higher abundances of methanotrophic taxa, indicating a direct microbial contribution to observed fluxes. For foliar N_2_O exchange, we hypothesize that (iv) flux magnitudes will differ among species according to variation in foliar transpiration, soil N_2_O flux, and soil total nitrogen, reflecting the integrated influence of physiological regulation and substrate availability rather than microbial mediation, with early-successional (pioneer) species exhibiting higher N_2_O emissions than late-successional species. By integrating in-situ flux measurements with leaf trait data, soil parameters, and microbial characterization for CH_4_, this study aims to elucidate the mechanistic controls of foliar GHG exchange and refine understanding of the role of temperate forest vegetation in ecosystem-scale CH_4_ and N_2_O budgets.

## METHODS

### Description of the study area

Field measurements were conducted at Downsview Park, a 260.6-ha urban green space in north-central Toronto, Ontario (43.7437° N, 79.4831° W), situated within the Humber River– Black Creek watershed and managed by Canada Lands Company (SM 1). The park lies within the Mixedwood Plains Ecozone (Genco, 2007; Crins *et al*., 2009). Our study focused on a mixed-species forest restoration site, where tree planting was undertaken during 2017–2018 with spatially interspersed species (SM 1). The landscape consists of gently undulating glacial till at 185–200 m elevation (Saurette *et al*., 2021) with generally low nutrient availability (Karim *et al*., 2025). The trial site was originally underlain by Jeddo clay loam, a poorly drained gleysol formed from glacial deposits (Saurette *et al*., 2021). Subsequent development has altered these soils into human-mixed and amended surface layers (De Sousa & Spiess, 2018). In the World Reference Base for Soil Resources, these are classified as “transportic” or “anthropic” (FAO, 2014; Burghardt *et al*., 2015), while the USDA system categorizes them as Human-Altered or Human-Transported (HAHT) or Human-Altered Material (HAM) (Galbraith, 2018). The disturbed soils and bulk density (∼1.22 g cm^−3^) fall within the range typical of urban industrial and parkland environments (Pouyat *et al*., 2007).

The regional climate is humid continental (Dfb), with a mean annual temperature of ∼9.7 °C and mean annual precipitation of ∼822 mm (Environment Canada, 2025). The herbaceous layer includes naturalized and native species such as *Solidago canadensis* and *Poa pratensis*. The park also supports a diverse assemblage of urban wildlife, including over 200 bird species– such as *Chaetura pelagica* (chimney swift), *Melanerpes carolinus* (red-bellied woodpecker), and *Colaptes auratus* (northern flicker)–as well as small mammals and pollinator habitats (Downsview Park, 2025). Collectively, the site represents a multifunctional urban forest ecotone.

### Species selection

A total of 25 tree species were selected for foliar gas-exchange measurements within the study site (Table 1). Species selection was based on two primary criteria: (i) accessibility of foliage within the measurement height range of 1.0–1.5□m above ground, suitable for chamber-based sampling, and (ii) representation across major functional groups, including deciduous broadleaf (n = 20) and coniferous needleleaf (n = 5) taxa. All selected species were sufficiently replicated within the study area, with a minimum of five individuals per species (n = 5) to support statistical comparisons. To incorporate key functional attributes potentially influencing foliar GHG exchange, species-specific tolerance values to shade, drought, and waterlogging were compiled from Niinemets & Valladares, (2006). These tolerance rankings are derived from a globally harmonized five-level scale (1 = very intolerant to 5 = very tolerant), constructed through cross-calibration of continental datasets for 806 temperate Northern Hemisphere woody species based on physiological and ecological evidence of growth under limiting light and water conditions.

**Table 1.**
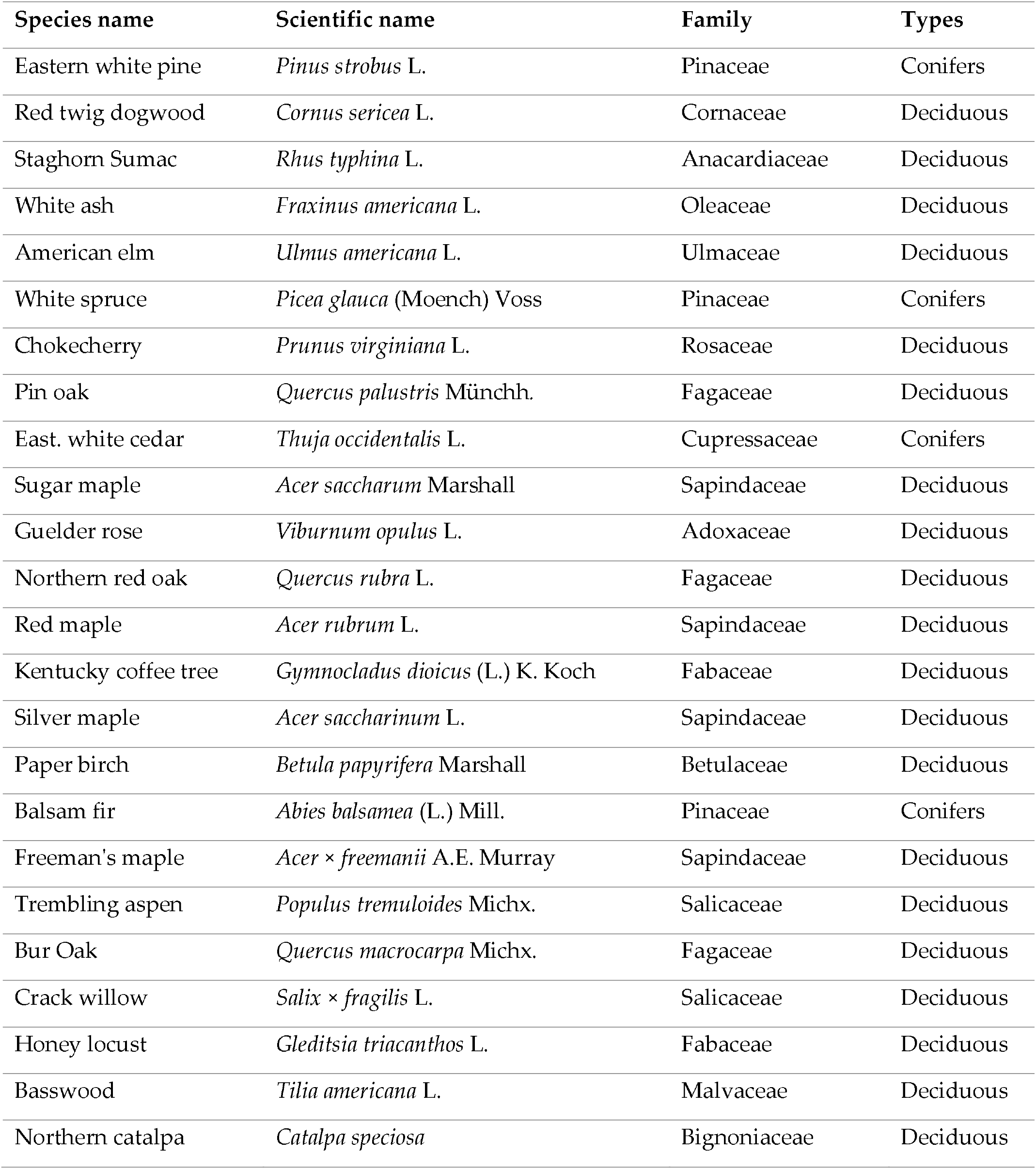
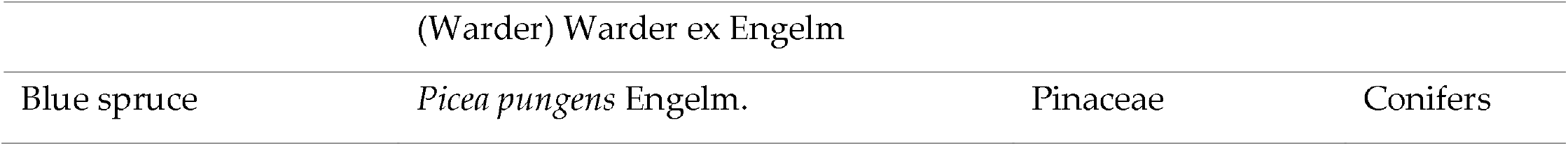
Species included in this study. Scientific names were verified and standardized using the NCBI Taxonomy database (Schoch *et al*., 2020).

| Species name | Scientific name | Family | Types |
| --- | --- | --- | --- |
| Eastern white pine | *Pinus strobus* L. | Pinaceae | Conifers |
| Red twig dogwood | *Cornus sericea* L. | Cornaceae | Deciduous |
| Staghorn Sumac | *Rhus typhina* L. | Anacardiaceae | Deciduous |
| White ash | *Fraxinus americana* L. | Oleaceae | Deciduous |
| American elm | *Ulmus americana* L. | Ulmaceae | Deciduous |
| White spruce | *Picea glauca* (Moench) Voss | Pinaceae | Conifers |
| Chokecherry | *Prunus virginiana* L. | Rosaceae | Deciduous |
| Pin oak | *Quercus palustris* Münchh. | Fagaceae | Deciduous |
| East. white cedar | *Thuja occidentalis* L. | Cupressaceae | Conifers |
| Sugar maple | *Acer saccharum* Marshall | Sapindaceae | Deciduous |
| Guelder rose | *Viburnum opulus* L. | Adoxaceae | Deciduous |
| Northern red oak | *Quercus rubra* L. | Fagaceae | Deciduous |
| Red maple | *Acer rubrum* L. | Sapindaceae | Deciduous |
| Kentucky coffee tree | *Gymnocladus dioicus* (L.) K. Koch | Fabaceae | Deciduous |
| Silver maple | *Acer saccharinum* L. | Sapindaceae | Deciduous |
| Paper birch | *Betula papyrifera* Marshall | Betulaceae | Deciduous |
| Balsam fir | *Abies balsamea* (L.) Mill. | Pinaceae | Conifers |
| Freeman's maple | *Acer × freemanii* A.E. Murray | Sapindaceae | Deciduous |
| Trembling aspen | *Populus tremuloides* Michx. | Salicaceae | Deciduous |
| Bur Oak | *Quercus macrocarpa* Michx. | Fagaceae | Deciduous |
| Crack willow | *Salix × fragilis* L. | Salicaceae | Deciduous |
| Honey locust | *Gleditsia triacanthos* L. | Fabaceae | Deciduous |
| Basswood | *Tilia americana* L. | Malvaceae | Deciduous |
| Northern catalpa | *Catalpa speciosa* | Bignoniaceae | Deciduous |

### Data collection

Our sampling strategy was designed to maximize both taxonomic breadth and within-species replication. Measurements were conducted during two distinct seasonal periods–early spring (mid-March 2024) and late autumn (mid-October 2024) –to capture phenological extremes in leaf development. The spring campaign coincided with the emergence of new leaves in deciduous broadleaf species, while the autumn campaign targeted fully mature or senescing foliage. In each season, foliar gas-exchange measurements were performed on 250 leaves from 125 individual trees, representing 25 (20 broadleaf deciduous and 5 needleleaf) species spanning 15 angiosperm and gymnosperm families.

Trees were selected from upland, non-inundated microsites across the study area, with spatial interspersion of individuals of different species. All sampled trees had a diameter at breast height (DBH) >1□cm (mean ± SD: 10.7□±□4.2□cm). For each species, gas-exchange measurements were made on two leaves per individual from five replicate trees (n = 10 leaves per species per season), targeting recently fully expanded foliage (Thomas & Bazzaz, 1999). All measurements were conducted under high light conditions, with leaves exposed to 1000□μmol·m^−2^·s^−1^ photosynthetic photon flux density (PPFD) to simulate natural mid-day irradiance on clear days. This irradiance level is consistent with standard protocols for light-saturated photosynthetic measurements in sun-acclimated leaves. Each measurement lasted 5 minutes, with three 1-minute slope intervals extracted to calculate foliar fluxes of CH_4_ and N_2_O, ensuring robust quantification.

Paired soil CH_4_ and N_2_O fluxes were measured within the rooting zone (∼2.5□m radius) of each sampled tree using a closed-dynamic chamber system (LI-8100A, LI-COR Inc., Lincoln, NE, USA) coupled to high-sensitivity analyzers (SM 2), following the configuration described in Karim et al., (2024). PVC collars (10□cm diameter) were inserted ∼3□cm into the soil at least 24 hours prior to flux measurements. Each soil flux measurement lasted 2.5 minutes and was replicated twice; mean values were used in subsequent analyses (Gorgolewski *et al*., 2022).

In addition to gas-exchange measurements, each measured leaf was collected for trait analyses. Specific leaf area (SLA) was calculated as the ratio of fresh leaf area (measured with an LI-3100C leaf area meter; LI-COR Inc., Lincoln, NE, USA) to oven-dried mass (Garnier *et al*., 2001). Foliar nitrogen concentration (%) was determined on dried samples using elemental analysis (CN628 Elemental Analyzer, LECO Co., St. Joseph, MI, USA). Tree DBH was recorded for all individuals. Concurrently, adjacent soil samples from the gas-flux collars were collected for analysis of total soil nitrogen concentration (%), providing an environmental context for microbial and physiological flux drivers.

Traditional static chamber methods combined with syringe sampling and gas chromatography have historically been used to measure CH_4_ and N_2_O fluxes from plant and soil systems (Machacova *et al*., 2021; Yang, 2022). However, these methods lack the sensitivity and temporal resolution required for detecting low-magnitude fluxes under in situ field conditions. Recent developments in laser-based spectroscopy have enabled high-frequency, sub-ppb measurements of trace gases with greater precision and reliability (Gorgolewski *et al*., 2023).

We employed a fully integrated system consisting of an off-axis integrated cavity output spectroscopy (OA-ICOS) analyzer for CO_2_, CH_4_, and H_2_O (LGR 915–0011; Los Gatos Research, CA, USA) and an optical feedback cavity-enhanced absorption spectroscopy (OF-CEAS) analyzer for N_2_O (LI-7820; LI-COR Inc., Lincoln, NE, USA). This combination has been recently validated for detecting low-level CH_4_ and N_2_O fluxes from tree foliage under field conditions (Karim *et al*., 2024, 2025), offering a robust and field-appropriate alternative to conventional laboratory-based approaches.

To minimize measurement artifacts, we used a dynamic flow-through leaf chamber (CS-LC7000; CredoSense Inc., Toronto, Canada) specifically designed for intact-leaf flux measurements under ambient field conditions. The chamber maintains a stable microenvironment with controlled airflow, temperature, and humidity (Karim *et al*., 2025), thereby reducing boundary layer resistance and preserving physiological leaf function (Ainsworth & Rogers, 2007; Engineer *et al*., 2016). Prior approaches using static chambers or modified soil collars have introduced significant uncertainties due to uncontrolled chamber atmospheres and variable effects of leaf detachment (Santiago & Mulkey, 2003; Gorgolewski *et al*., 2023).

Each measurement consisted of a 5-minute cycle, comprising three closed-dynamic loops (60 seconds each) interspersed with 60-second ambient air flushing phases, regulated by an automated valve system (SM 2). This design prevents trace gas accumulation and maintains ambient CO_2_ concentration, ensuring consistent stomatal conductance throughout the measurement period. The chamber was equipped with a full-spectrum photodiode light source delivering 1000□μmol·m^−2^·s^−1^ of photosynthetically active radiation (PAR). Leaves were acclimated to this irradiance for at least 5 minutes to reach steady-state gas exchange.

### Flux calculations

Concentration data for CO_2_, CH_4_, H_2_O, and N_2_O were collected using two analyzers and merged by timestamp after conversion to a common unit (ppm). To minimize measurement artifacts, the initial 10□s of each 1-minute leaf chamber closure and the initial 15□s of each 2-minute soil chamber closure were excluded as “dead band” intervals (Hoffmann *et al*., 2017).

Flux calculation time windows were selected based on CO_2_ data, which tend to exhibit higher signal-to-noise ratios and are more sensitive to leakage or pressure inconsistencies than CH_4_ or N_2_O (Karim *et al*., 2024; Halim *et al*., 2024). A moving window (35–50□s for leaves; 60– 105□s for soils) was evaluated using the Pearson correlation coefficient (*r*) between concentration and time. The window with the highest *r*-value was chosen for slope computation and applied consistently to all gases in the same chamber run.

To calculate the slope (*dc/dt*) for CO_2_, CH_4_, H_2_O, and N_2_O fluxes, we utilized either linear or non-linear regression, following (Halim *et al*., 2024). If the quadratic term in a polynomial fit was non-significant (*p* > 0.05), we used a linear fit, otherwise we choose a non-linear fit. Flux (F) was then computed using the following equation 1 (LI-COR, 2015),

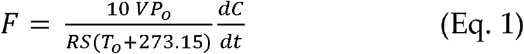

Where F is the gas flux, V is the chamber headspace volume (cm^3^), P_o_ is the initial pressure (kPa), R is the universal gas constant (0.83144598 m^3^·kPa·K^−1^·mol^−1^), To is the initial temperature (K), S is the projected surface area of the leaf or soil (cm^2^), and *dC/dt* is the initial slope of concentration change (μmol·mol^−1^·s^−1^). Throughout this paper, CO_2_ fluxes are reported as μmol·mol^-1^·s^-1^, CH_4_ fluxes as nmol·m^-2^·s^-1^, H_2_O fluxes as mmol·m^-2^·s^-1^, and N_2_O fluxes as pmol·m^-2^·s^-1^. All fluxes are expressed per unit area of the measured surface–soil fluxes per unit soil surface area, and foliar fluxes per unit leaf surface area.

For cases where the quadratic coefficient was significant (*p*□<□0.05), a nonlinear model was applied by fitting the following empirical equation 2 (Welles *et al*., 2001):

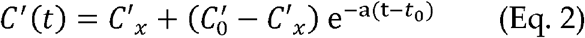

Where *C′(t)* is the instantaneous H_2_O, and water-corrected CO_2_ or CH_4_ or N_2_O mole fraction, *C*_*o*_′ is the value of *C′(t)* when the chamber just closed, *C*_*x*_′ is the asymptote parameter, *a* specifies the curvature of the fit (*s*^*-1*^), and *t*_*o*_ is time (*s*) when the chamber closed. The parameters *a, t*_*o*_, *C*_*x*_′, and *C*_*o*_′ were estimated from the fitted nonlinear regression. Subsequently, using the following equation (Eq. 3), which is derived from Eq. 2 (at *t* = *t*_*o*_) was used to calculate the initial rate of change of the H_2_O, and water-corrected CO_2_, CH_4_, or N_2_O) mole fraction (LI-COR, 2015):

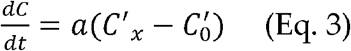

The resulting *dc/dt* value was then inserted into in Eq. 1 to obtain gas fluxes. Overall, using the above algorithm, 8% of CO_2_ fluxes, 11% of CH_4_ fluxes, 6% of H_2_O fluxes, and 13% of N_2_O fluxes required a non-linear fit, predominantly corresponding to high-flux measurements. We then averaged three replicate flux measurements per leaf and two replicate flux measurements per soil collar for further analyses.

### Leaf and soil total N measurement

Leaf and soil samples were dried at 60°C for 12 hours and then finely ground (< 0.5 mm). Prior to combustion, the ground samples were further dried for 30 minutes at 60°C to remove any residual moisture. Each sample (20 g) was weighted before and after combustion to assess the loss of organic matter. Total nitrogen (N) content was determined using a LECO 628 Series CN analyzer (LECO Corporation, St. Joseph, MI, USA). During high-temperature combustion in an O_2_-rich atmosphere, nitrogen was converted to nitrogen oxides (NO_*x*_), which were subsequently quantified. Instrument calibration was performed using Elemental Drift Reference (EDR) standards.

### Sample collection and processing for leaf microbial analysis

For evidence of leaf-associated methanotrophs and detailed taxonomic/functional characterization, we selected two focal tree species for representing the extremes of measured foliar CH_4_ oxidation: *T. americana* (highest CH_4_ oxidation) and *P. virginiana* (lowest CH_4_ oxidation). From each species we collected three biological replicates (n = 3 individuals per species; total n = 6) on Sept. 11, 2025. Replicates were chosen to match the individuals used for flux measurements in all cases, and sampling targeted the same crown exposure and height used during flux measurement to minimize microenvironmental differences. From each selected tree, we excised three fully expanded, healthy leaves from the same aspect and height used for the gas flux measurement. Leaves were handled using sterile nitrile gloves and sterilized scissors; each pooled sample consisted of leaves representing ∼20–50 cm^2^ total surface area (recorded per sample for normalization). Immediately after excision, leaves were placed into sterile 50-mL Falcon tubes (dry, without buffer) and kept on ice in a dark cooler for transport to the sequencing lab (Metagenom Bio Life Science Inc., Waterloo, ON, Canada) for DNA extraction and sequencing.

### Microbial analysis and data processing, cleaning and normalization

Genomic DNA was extracted from pooled leaf samples using the Sox DNA Isolation Kit (Metagenom Bio Life Science Inc., Waterloo, ON, Canada) according to the manufacturer’s instructions. To characterize bacterial communities, with particular focus on methanotrophs, the V4 region of the 16S rRNA gene was amplified using PCR. Each reaction was performed in triplicate with a total volume of 25 μL per reaction. The reaction mixture contained 2.5 μL of 10× standard Taq buffer, 0.5 μL of 10 mM dNTPs, 0.25 μL of bovine serum albumin (20 mg/mL), 5 μL of 1 μM forward primer 515FB (5’-GTGYCAGCMGCCGCGGTAA) and 5 μL of 1 μM reverse primer 806RB (5’-GGACTACNVGGGTWTCTAAT) (William *et al*., 2015), 5 μL of DNA template, 0.2 μL of Taq DNA polymerase (5 U/μL), and 6.55 μL of PCR-grade water. Thermal cycling consisted of an initial denaturation at 95°C for five minutes, followed by 35 cycles of denaturation at 95°C for 30 seconds, annealing at 30°C for 30 seconds, and extension at 72°C for 50 seconds, with a final extension at 72°C for ten minutes. Triplicate PCR products were pooled and visualized on a 2% TAE agarose gel to confirm amplicons of the expected size. Correct amplicons were gel-purified, quantified using the Qubit dsDNA HS Assay Kit (Thermo Fisher Scientific Inc., Massachusetts, USA), and pooled in equimolar concentrations for sequencing. Libraries were sequenced on the Illumina MiSeq platform using the MiSeq Reagent Kit v2 (2 × 250 cycles).

Demultiplexed sequences were processed using *cutadapt* for primer removal (Martin, 2011), followed by quality filtering, denoising, chimera removal, and amplicon sequence variant (ASV) inference using DADA2 v1.22 (Callahan *et al*., 2016). Taxonomy was assigned to representative sequences using *‘IdTaxa’* from the DECIPHER package in R (v 4.4.1) (Murali *et al*., 2018) trained against SILVA release 138 (16S rRNA; Quast et al., (2013)). Non-target sequences, including chloroplast and mitochondrial reads, were removed prior to downstream analyses. Samples were rarefied to the sequence count of the smallest library exceeding 5,000 reads to standardize sequencing depth. Functional characterization of methanotrophs via pmoA analysis was also conducted.

Overall, the sequencing quality was high, with an average read recovery of 92.3 ± 1.4% for the 16S dataset and up to 73% for pmoA, indicating robust data retention and minimal chimera formation. Detailed read processing statistics for each sample are provided in SM 3.

### Statistical analysis

All flux calculations and statistical analyses were performed in R (version 4.4.2) (R Core Team, 2024). To assess the effects of season and species type on foliar CH_4_ and N_2_O fluxes, linear mixed-effects models were fitted using the *lmer* function from the *lme4* package (Bates *et al*., 2015). Season (spring, fall) and shade-tolerance class were included as categorical fixed effects, along with their interaction (season × shade-tolerance class). Tree identity was included as a random effect to account for repeated measurements within individual trees. Fixed effects were tested using Two-way ANOVA with Satterthwaite’s approximation for degrees of freedom, ensuring independent evaluation of main effects and their interaction in this factorial design. Model assumptions were assessed using quantile–quantile plots and Shapiro–Wilk tests for normality *(shapiro*.*test)* and Bartlett’s test for homogeneity of variances *(bartlett*.*test)*. Analysis of variance (ANOVA) was performed using *aov* from the base *stats* package, followed by Tukey’s HSD post-hoc comparisons (TukeyHSD). Two-sample t-tests *(t*.*test)* were applied to compare mean fluxes within each species between spring and fall seasons. To partition variance hierarchically among families, species within families, individual trees, and measured leaves, nested models were fitted using the ‘*lme’* function in the *nlme* package (Pinheiro *et al*., 2024). Detection limits for foliar CH_4_ and N_2_O fluxes were estimated from the minimum statistically significant slopes in gas concentration versus time across all empirical measurements, reflecting the sensitivity of the instrumentation and protocol.

To complement the taxonomy-based analysis, a species-level phylogenetic signal assessment was performed. A phylogenetic tree for the 25 species in the dataset was constructed using *Phylot v2*, a tree generator based on the NCBI taxonomy database (Schoch *et al*., 2020). The resulting ‘newick’ tree incorporated branch lengths corresponding to clade ages derived from fossil records (Wikström *et al*., 2001), as updated by (Gastauer & Meira-Neto, 2016). Using this phylogeny, Pagel’s λ was calculated for leaf CH_4_ and N_2_O fluxes with the ‘*phytools’* R package (Revell, 2024). Pagel’s λ serves as a scaling parameter for the phylogeny, quantifying the extent to which trait variation reflects evolutionary relatedness (Pagel, 1999). Values of λ near 1 indicate strong phylogenetic dependence, whereas values near 0 indicate that fluxes are distributed independently of phylogeny. The robustness of the λ estimates was evaluated via bootstrapped 95% confidence intervals. The phylogenetic tree and associated trait data were visualized using the *‘ggtree’* function in R (Yu *et al*., 2017).

## RESULTS

Foliar CH_4_ oxidation and N_2_O emission showed distinct patterns. All measured leaves acted as a net CH_4_ sink, with mean uptake of 1.03 ± 0.04 nmol·m^−2^·s^−1^, while N_2_O fluxes were uniformly positive, with a mean emission of 1.01 ± 0.047 pmol·m^−2^·s^−1^. CO_2_ and H_2_O fluxes confirmed active leaf physiology: leaves maintained steady CO_2_ uptake (−11.6 ± 0.25 μmol·m^−2^·s^−1^) and released H_2_O (2.58 ± 0.068 mmol·m^−2^·s^−1^) through transpiration. Physiological activity remained strong throughout the study period. Species identity contributed substantially to variation in CO_2_ uptake (F = 33.07, *p* < 0.001) and H_2_O flux (F = 11.02, *p* < 0.001), with negligible seasonal effects (Table 2).

**Table 2.**
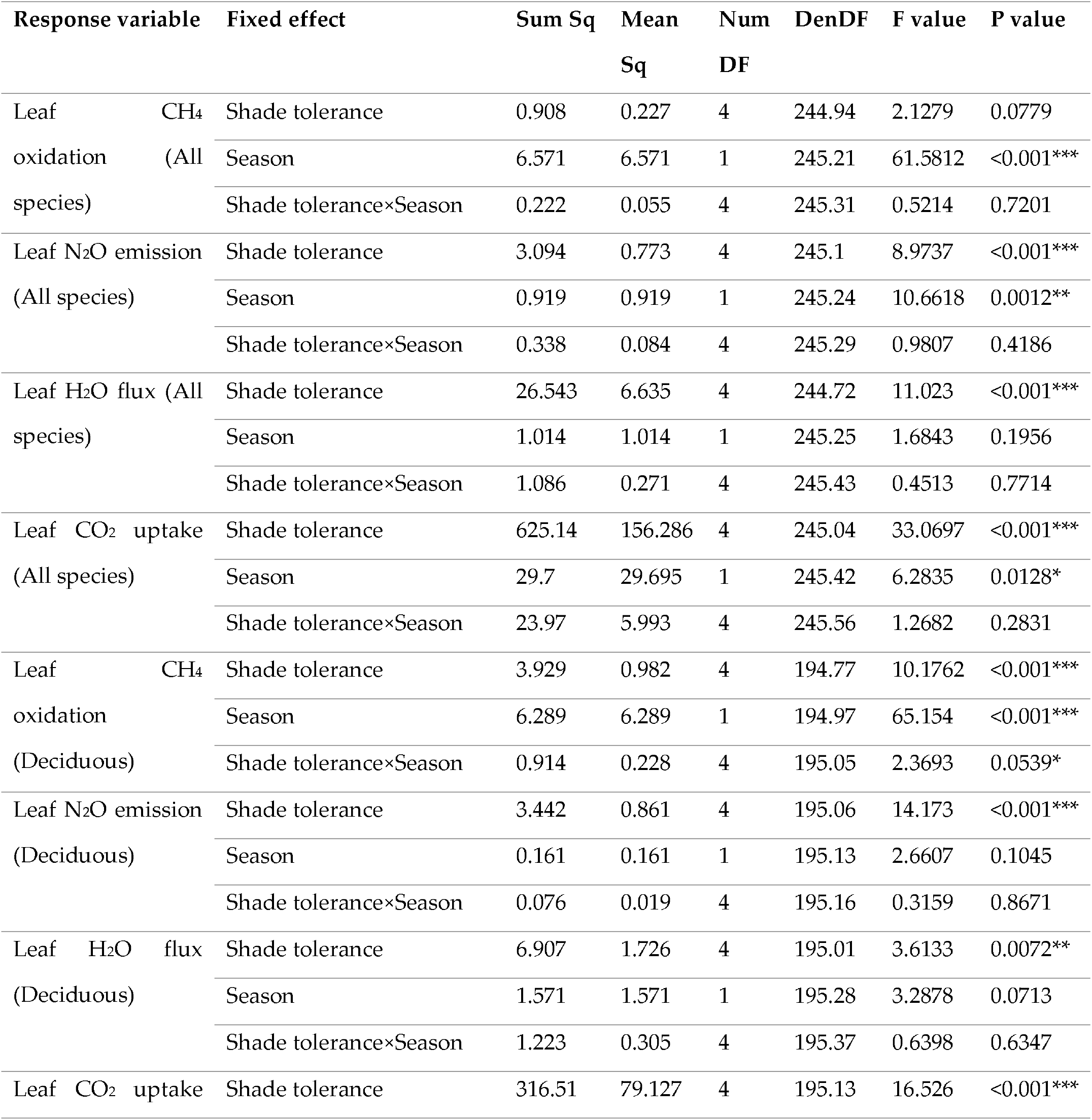
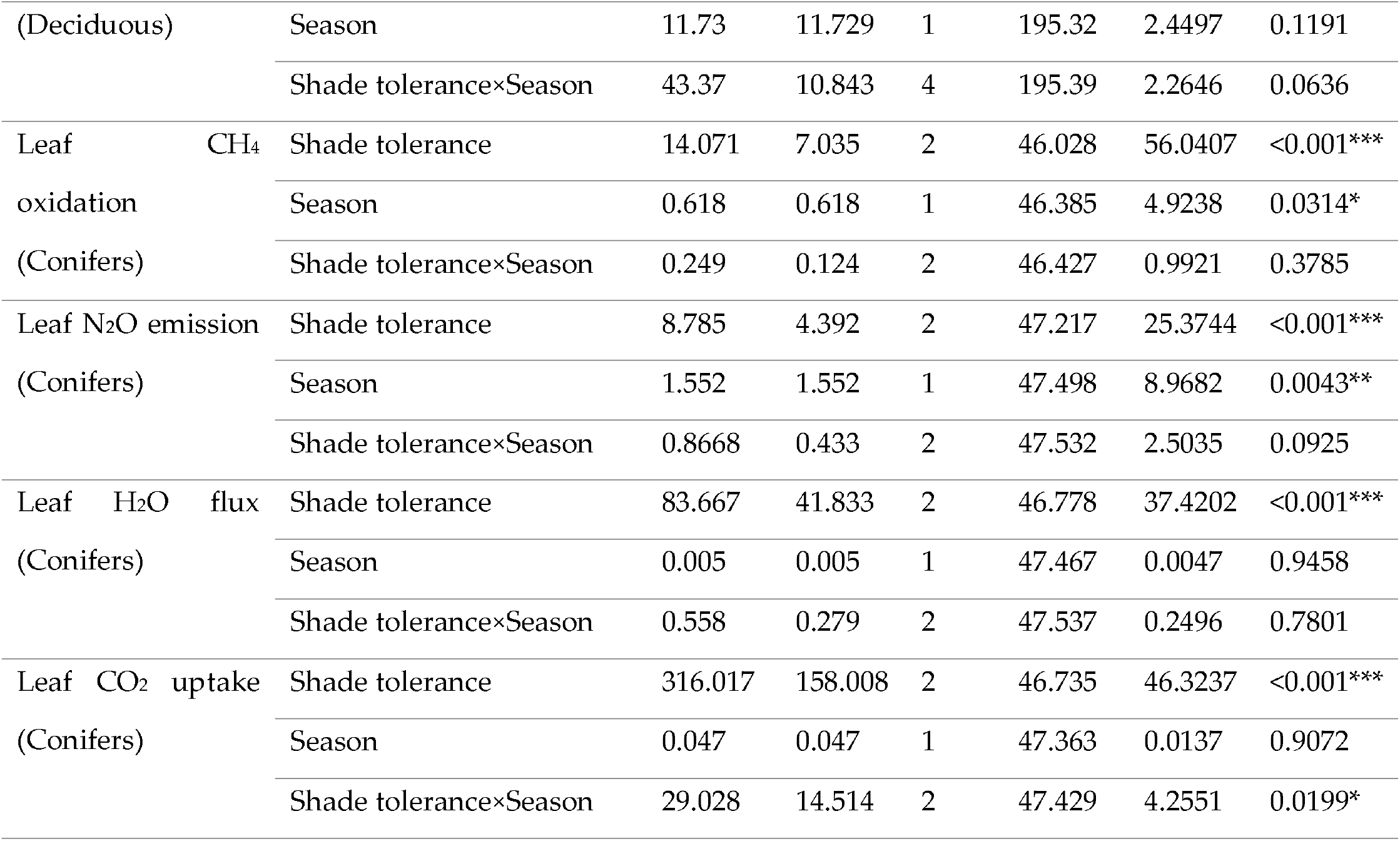
Seasonal and Shade tolerance class effects on foliar CO_2_, CH_4_, N_2_O, and H_2_O fluxes, based on linear mixed-effects models. ANOVA statistics (df, F-value, and p-value) indicate the significance of main and interactive effects between species type (deciduous vs. coniferous and overall) and season (spring 2024 vs. fall 2024).

### Foliar CH_4_ and N_2_O flux variation with shade tolerance and season

Foliar CH_4_ oxidation varied significantly with shade tolerance and season (Table 2).

In deciduous (angiosperm) species, CH_4_ oxidation increased with shade-tolerance class. Shade-intolerant species (class 1) had the lowest uptake, averaging 0.497 ± 0.10 nmol·m^−2^·s^−1^ in spring and 0.754 ± 0.14 nmol·m^−2^·s^−1^ in fall (SM 4), whereas shade-tolerant species (class 5) reached 1.80 ± 0.17 nmol·m^−2^·s^−1^ in spring and 2.05 ± 0.16 nmol·m^−2^·s^−1^ in fall (Figure 1a, SM 4). Seasonal variation amplified this trend in shade-tolerant to mid-tolerant deciduous trees, with shade-tolerance classes 2–4 showing significant increases from spring to fall (pairwise t-tests, *p* < 0.05).

**Figure 1.**
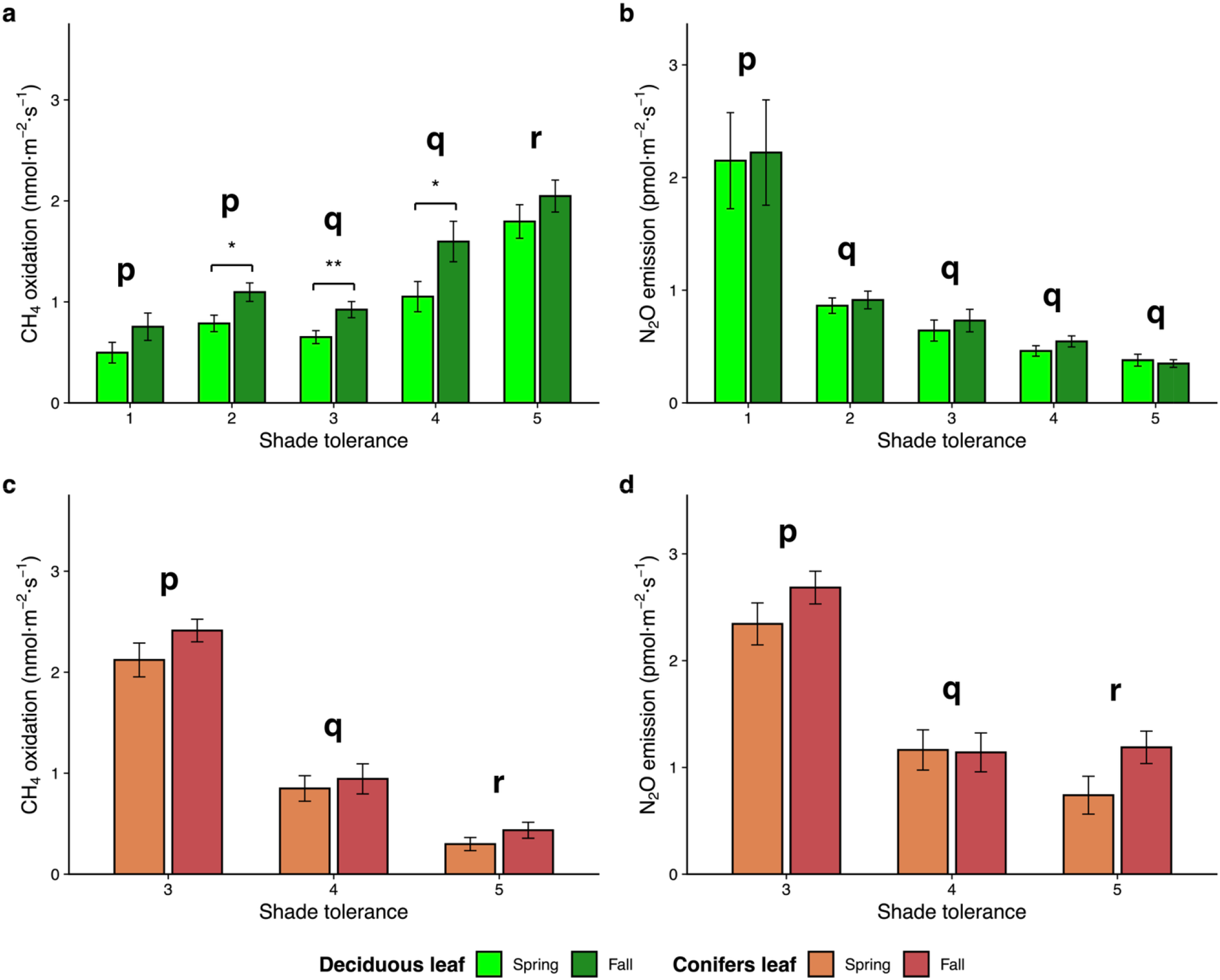
Effects of shade tolerance on foliar CH_4_ oxidation and N_2_O emission in deciduous and conifer leaves. Bars show mean fluxes (± SE) for each shade-tolerance class, with Spring and Fall displayed separately. Lowercase letters indicate significant differences among shade-tolerance classes (Tukey’s HSD, *p* < 0.05). Asterisks positioned between seasonal bars denote significant differences between spring and fall within each shade-tolerance class based on pairwise t-tests (\**p* < 0.05, \*\**p* < 0.01, \*\*\**p* < 0.001).

Conifers exhibited a different pattern: mid-tolerant species (class 3) showed the highest uptake among all groups, 2.12 ± 0.17 nmol·m^−2^·s^−1^ in spring and 2.41 ± 0.11 nmol·m^−2^·s^−1^ in fall (SM 4). Shade-tolerant conifers (shade tolerance class 5) declined sharply, averaging 0.298 ± 0.07 nmol·m^−2^·s^−1^ in spring and 0.435 ± 0.08 nmol·m^−2^·s^−1^ in fall (Figure 1c, SM 4). Seasonal differences in conifers were minimal and not statistically significant (*p > 0*.*05*). Foliar CH_4_ uptake was higher in conifers than deciduous species in mid-tolerant species (SM 5, SM 6), but this pattern was reversed among shade-tolerant species (Figure 1, SM 5, SM 6).

Foliar N_2_O fluxes showed the opposite trend, with shade-intolerant and mid-tolerant species being the main emitters. In deciduous trees, intolerant species (class 1) released 2.15 ± 0.43 pmol·m^−2^·s^−1^ in spring and 2.22 ± 0.47 pmol·m^−2^·s^−1^ in fall (Figure 1b, SM 4), substantially exceeding mid-tolerant and shade-tolerant classes (Tukey HSD, *p* < 0.001), which ranged from 0.35–0.90 pmol·m^−2^·s^−1^. Conifers followed a similar pattern: mid-tolerant species (class 3) produced the highest fluxes (2.34 ± 0.20 pmol·m^−2^·s^−1^ in spring; 2.68 ± 0.15 pmol·m^−2^·s^−1^ in fall), while shade-tolerant conifers (class 5) emitted far less, averaging 0.74 ± 0.18 pmol·m^−2^·s^−1^ in spring (Figure 1d, SM 5). N_2_O fluxes in conifers were stable across seasons (all *p* > 0.05, SM 6).

Drought tolerance affected both gases (*p* < 0.001): the most drought-tolerant species had lowest CH_4_ oxidation and highest N_2_O emissions (*p* < 0.05) (SM 4, SM 7, SM 8A, SM 8B). Waterlogging tolerance also influenced both CH_4_ oxidation (*p* < 0.001) and N_2_O emission (*p* < 0.01) (SM 4, SM 7, SM 8C, SM 8D), with least tolerant species showing the highest CH_4_ uptake and most tolerant species showing lowest N_2_O release.

### Taxonomic variation and phylogenetic structure of foliar CH_4_ and N_2_O fluxes

Foliar CH_4_ oxidation and N_2_O emission varied markedly across species and families, reflecting the influence of taxonomic identity on leaf-level GHG fluxes. CH_4_ oxidation among the 25 species ranged from near zero in *P. virginiana* (0.027–0.050□nmol·m^−2^·s^−1^) to consistently high uptake in *T. americana* (2.09–3.01), *T. occidentalis* (2.65–2.74), and *C. speciosa* (1.95– 2.20□nmol·m^−2^·s^−1^). Several deciduous hardwoods (*A. rubrum, A. saccharum, Q. rubra* >1.5□nmol·m^−2^·s^−1^) and the conifer *P. strobus* also maintained elevated oxidation (Figure 2a). Seasonal contrasts, confirmed by Tukey’s post-hoc tests, generally showed higher oxidation in fall, with significant differences in *A. freemanii* (*p*□<□0.001), *A. saccharinum* (*p*□=□0.03), *B. papyrifera* (*p*□=□0.003), *C. sericea* (*p*□<□0.001), *F. americana* (*p*□<□0.001), *P. strobus* (*p*□=□0.038), *P. tremuloides* (*p*□<□0.001), *P. virginiana* (*p*□=□0.048), *Q. macrocarpa* (*p*□=□0.004), *Q. rubra* (*p*□<□0.001), *S. fragilis* (*p*□<□0.01), *T. americana* (*p*□<□0.001), and *V. opulus* (*p*□=□0.019). At the family level, CH_4_ oxidation mirrored these trends, ranging from 0.027□nmol·m^−2^·s^−1^ in Rosaceae to 3.01□nmol·m^−2^·s^−1^ in Malvaceae (SM 9a).

**Figure 2.**
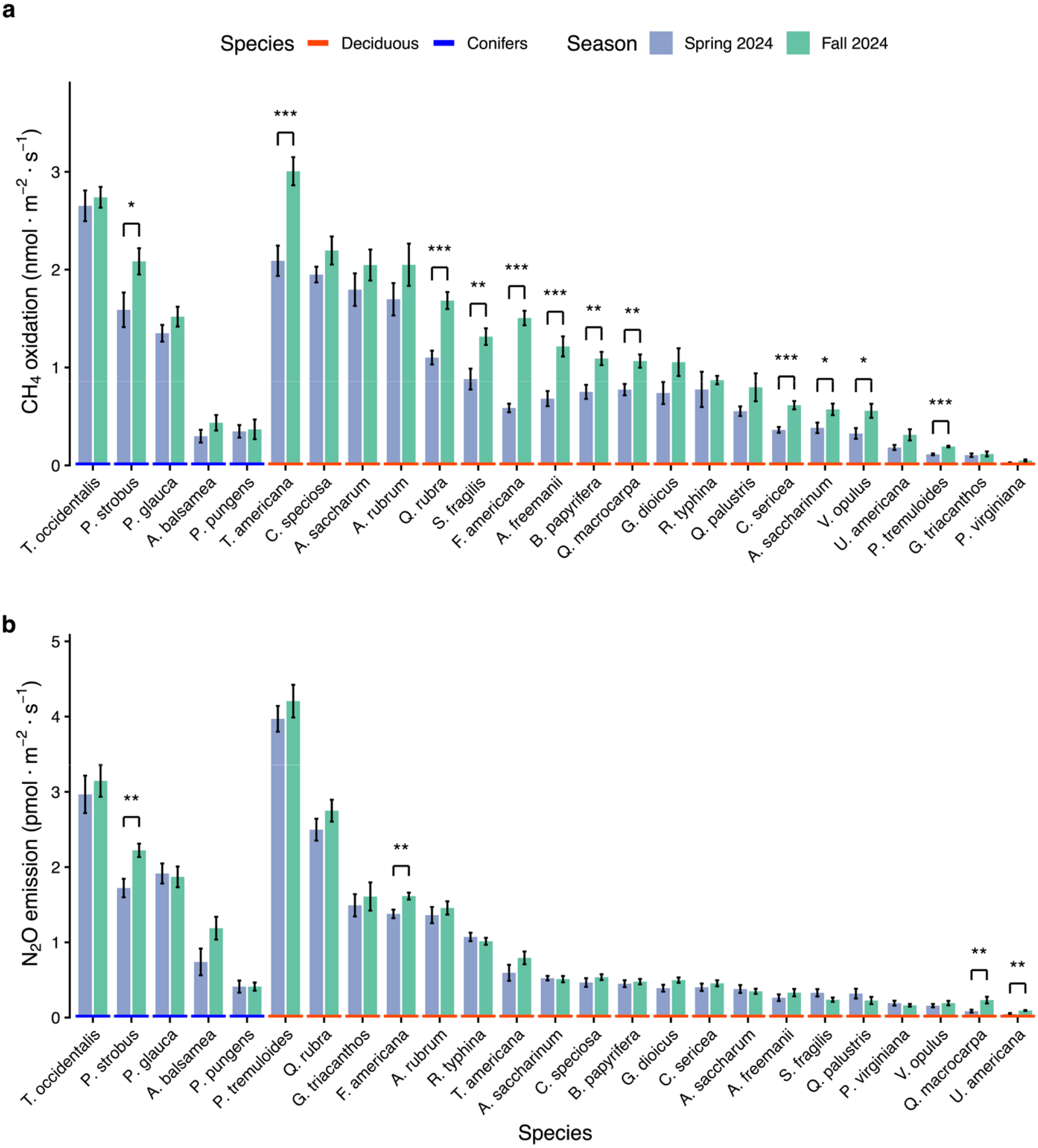
Seasonal variation in foliar GHG fluxes across 25 tree species. (a) Mean (± SE) CH_4_ oxidation (nmol·m^−2^·s^−1^) and (b) mean (± SE) N_2_O emission (pmol·m^−2^·s^−1^) measured on individual leaves during spring (green) and fall (blue) 2024. Species are ordered within deciduous and conifer groups by decreasing mean flux. Colored baselines indicate functional type (orange for deciduous, blue for conifers). Asterisks denote significant seasonal differences (\**p* < 0.05, \*\**p* < 0.01, \*\*\**p* < 0.001).

N_2_O emissions also varied among species (Figure 2b), from < 0.3□pmol·m^−2^·s^−1^ in *A. freemanii* and *A. saccharum* to higher rates in *A. rubrum* (1.36–1.46), *A. balsamea* (0.74–1.19), and *B. papyrifera* (0.92–1.11□pmol·m^−2^·s^−1^). Conifers, including *P. strobus* and *P. virginiana*, consistently showed low N_2_O fluxes. Seasonal variation in N_2_O emissions was generally less pronounced than for CH_4_ at the family level (SM 9b). These results reveal a clear taxonomic hierarchy in foliar GHG fluxes.

Phylogenetic analysis showed no detectable signal for either gas. Foliar CH_4_ oxidation had Pagel’s λ = 0.012 (p = 1, 95% CI = [0, 0.861]; log-likelihood −31.121 vs. −31.120 for λ = 0; SM 10a), while N_2_O emission had λ = 1 × 10^−4^ (p = 1, 95% CI = [0, 0.981]; log-likelihoods identical at −38.214; SM 10b). These findings indicate that variation in CH_4_ and N_2_O fluxes is independent of phylogeny, suggesting species-specific traits or environmental factors exert stronger influence than evolutionary history.

### Leaf fluxes in relation to soil fluxes and leaf traits

Foliar gas fluxes correlated with soil fluxes and leaf traits (Figure 3). Leaf CH_4_ oxidation significantly related with soil CH_4_ oxidation (R^2^ = 0.23, *p* < 0.001; Figure 3a), whereas leaf N_2_O emissions increased with soil N_2_O emissions (R^2^ = 0.26, *p* < 0.001; Figure 3b). Associations with leaf transpiration were weaker: CH_4_ oxidation showed a marginal positive trend (R^2^ = 0.02, *p* < 0.01; Figure 3c), whereas N_2_O emissions followed an increasing trend (R^2^ = 0.18, *p* < 0.001; Figure 3d). Leaf nitrogen content was weakly but significantly associated with CH_4_ oxidation (R^2^ = 0.03, *p* < 0.001; Figure 3e) and more strongly with N_2_O emissions (R^2^ = 0.11, *p* < 0.001; Figure 3f). Structural traits had comparatively minor effects: CH_4_ oxidation showed a weak response to SLA (R^2^ = 0.01, *p* < 0.05; Figure 3g), while N_2_O emissions declined linearly with SLA (R^2^ = 0.04, *p* < 0.001; Figure 3h). Overall, soil gas fluxes showed the strongest correlations with foliar CH_4_ oxidation and N_2_O emissions, with leaf traits contributing weaker associations.

**Figure 3.**
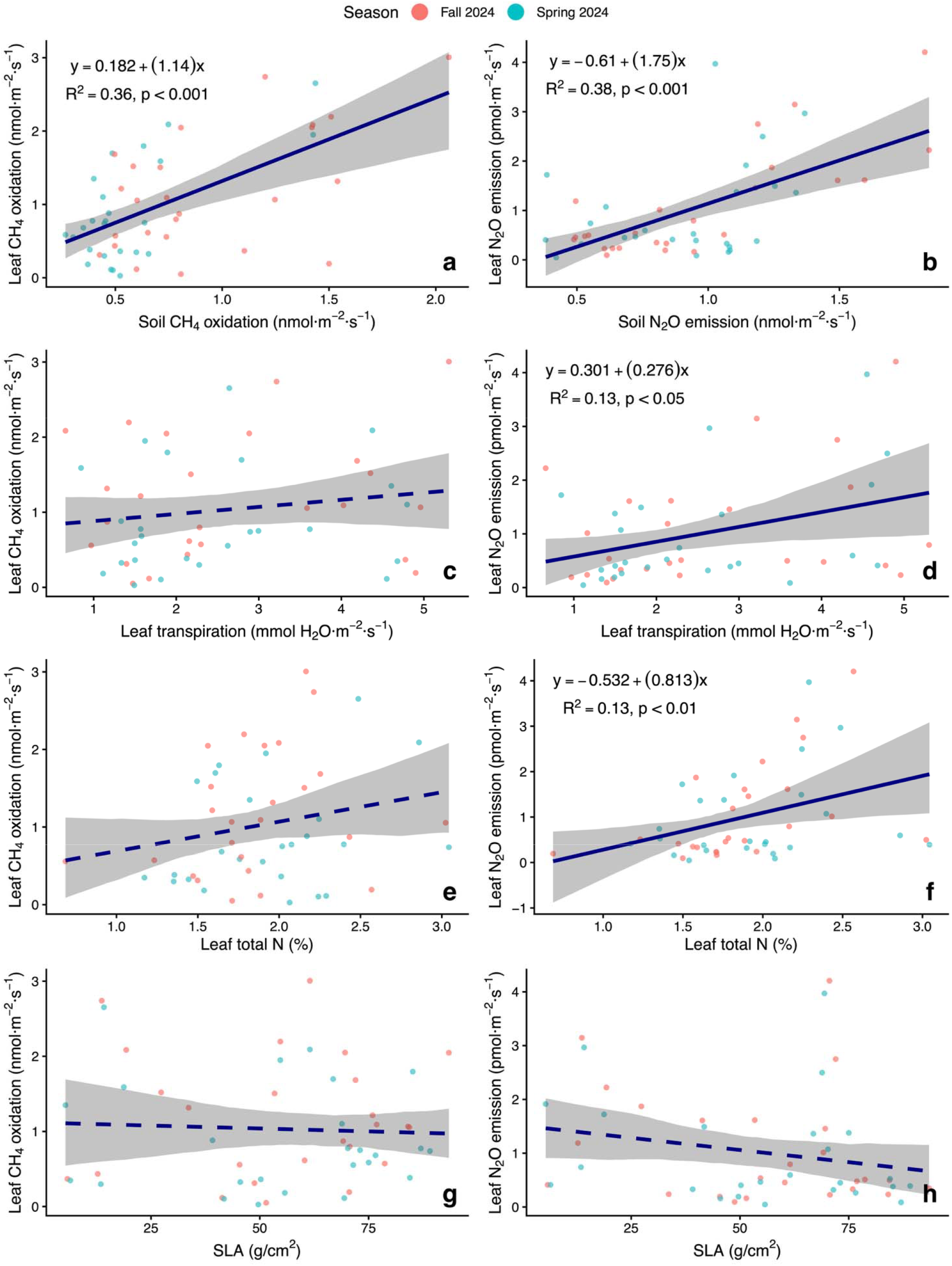
Relationships between foliar gas fluxes and biophysical or biogeochemical factors. Panels (a, c, e, g) show relationships between leaf CH_4_ oxidation rates (nmol·m^−2^·s^−1^) and (a) soil CH_4_ oxidation (nmol·m^−2^·s^−1^), (c) leaf transpiration (mmol H_2_O·m^−2^·s^−1^), (e) leaf total nitrogen (%), and (g) specific leaf area (SLA; g·cm^−2^). Panels (b, d, f, h) show relationships between leaf N_2_O emission rates (nmol·m^−2^·s^−1^) and (b) soil N_2_O emission (nmol·m^−2^·s^−1^), (d) leaf transpiration (mmol H_2_O·m^−2^·s^−1^), (f) leaf total nitrogen (%), and (h) SLA (g·cm^−2^). Data points are colored by season: Spring 2024 (blue) and Fall 2024 (orange). Dark blue lines represent best-fit regression models with 95% confidence intervals (shaded areas). Solid lines indicate significant relationships where *p*>0.05, while dashed lines indicate weak relationships where *p* < 0.05.

### Nested analysis of variance of foliar CH_4_ and N_2_O fluxes among hierarchical levels

For foliar CH_4_ oxidation, species within families accounted for the largest variance component (variance = 0.550; SD = 0.742; 80.0% of total; SM 11). Variability among individual trees within species was smaller (variance = 0.011; SD = 0.105; 1.6%), family-level variance was negligible (variance = 8.7 × 10^−10^; SD = 2.95 × 10^−5^; ≈ 0%), and leaf-level variance was minimal (2.7 × 10^−10^; SD = 1.64 × 10^−5^; ≈0 %). Residual variance was 0.124 (SD = 0.352; 18.0 %).

For foliar N_2_O fluxes, species-level variance was dominant (variance = 1.028; SD = 1.014; 91.0%). Variance among individual trees within species was small (variance = 5.57 × 10^−3^; SD = 0.075; 0.49%), family-level variance negligible (variance = 2.69 × 10^−8^; SD = 1.64 × 10^−4^; ≈ 0%), and leaf-level variance was 6.02 × 10^−3^ (SD = 0.078; 0.53%). Residual variance was 0.092 (SD = 0.303; 8.1%). Thus, species-level variation was the dominant source for both CH_4_ and N_2_O fluxes, with residual variance larger for N_2_O.

### Methanotrophic community composition between high and low CH_4_-uptake species

Amplicon sequencing of 16S rRNA and pmoA genes revealed pronounced contrasts in methanotrophic community structure between the high methane-uptake species *T. americana* and the low-uptake species *P. virginiana*. Methanotrophs were detected in all samples, but their taxonomic composition, relative abundance, and functional gene representation differed between host species (Table 3).

**Table 3.** Relative abundance (%) of dominant methanotrophic and associated taxa in replicate samples of *Tilia americana* (Basswood; high CH_4_ uptake) and *Prunus virginiana* (Chokecherry; low CH_4_ uptake). Abbreviations: Tilia = *T. americana*; Pru = *P. virginiana*.

| Genus | Family | Type | Tilia<br>01 | Tilia<br>02 | Tilia<br>03 | Pru<br>01 | Pru<br>02 | Pru<br>03 |
| --- | --- | --- | --- | --- | --- | --- | --- | --- |
| *Candidatus*<br>*Methylospira* | Methylococcaceae | Type I | 3.6 | 3.2 | 6.4 | 0.32 | 0.16 | 0.12 |
| *Methylomicrobium* | Methylomonadaceae | Type I | 0.4 | 0.7 | 0.3 | 0.01 | 0.0 | 0.0 |
| *Methylobacterium–*<br>*Methylorubrum* | Beijerinckiaceae | Facultative<br>Type II | 4.8 | 2.2 | 5.8 | 10.1 | 9.9 | 6.2 |
| *Methylocaldum* | Methylococcaceae | Type I | 0.6 | 0.2 | 0.09 | 0.0 | 0.0 | 0.0 |

In *T. americana*, the methanotrophic assemblage was dominated by obligate Type I methanotrophs (*Methylococcales*). Based on 16S rRNA data, *Candidatus Methylospira* (*Methylococcaceae*) constituted 3.2–6.4% of the bacterial community, the most abundant methane-oxidizing lineage. Additional Type I taxa, including *Methylomicrobium* (*Methylomonadaceae*; 0.3–0.7%) and *Methylocaldum* (*Methylococcaceae*; 0.09–0.6%), were consistently detected across replicates. pmoA sequencing yielded 48,298, 29,265, and 31,903 high-quality reads, approximately an order of magnitude higher than *P. virginiana*, with dominant ASVs aligning with particulate methane monooxygenase (pMMO) genes from *Methylococcales*.

In contrast, *P. virginiana* exhibited markedly lower Type I methanotroph abundance. *Candidatus Methylospira* and *Methylomicrobium* occurred at trace levels (< 0.4%), and *Methylocaldum* was absent. The community was enriched in facultative Type II methylotrophs *Methylobacterium–Methylorubrum* (*Beijerinckiaceae*; 6.2–10.1%). The pmoA sequencing produced 6,083, 981, and 104 reads, with dominant ASVs aligning with *Methylocystis* sp., a Type II methanotroph. Combined 16S rRNA and pmoA data indicate that *T. americana* hosts a Type I-dominated methanotrophic community with higher read counts, whereas *P. virginiana* harbors a functionally reduced assemblage dominated by facultative Type II and methylotrophic taxa.

### Species-level CH_4_ and N_2_O contributions to foliar GHG impact and restoration potential

Foliar GHG contributions were expressed as CO_2_-equivalents using GWP values of 27 for CH_4_ and 273 for N_2_O (Table 4). Across all leaves, CH_4_ oxidation dominated the GHG balance, with mean CH_4_-CO_2_ equivalents of 27.8□nmol·m^−2^·s^−1^ versus 0.28□nmol·m^−2^·s^−1^ for N_2_O, yielding an average CH_4_:N_2_O impact ratio of 258. Conifers showed moderate CH_4_ dominance (mean ratio 96), whereas deciduous species had higher relative CH_4_ impact (mean ratio 298). Species-level ratios ranged from 3.8 in *P. tremuloides* to 1,135 in *Q. macrocarpa* (SM 12). *Q. macrocarpa, A. saccharum*, and *U. americana* exhibited the highest CH_4_ uptake relative to N_2_O, while lower-ratio species contributed more evenly across both gases.

**Table 4.** Species-level GWP-weighted foliar CH_4_ uptake and N_2_O emission, ranked by the CH_4_-to-N_2_O impact ratio to identify species with the highest potential for climate-relevant restoration. The table includes mean GWP-weighted CH_4_ uptake (nmol CO_2_-eq·m^−2^·s^−1^), mean N_2_O emission (nmol CO_2_-eq·m^−2^·s^−1^), the CH_4_-to-N_2_O impact ratio, and a priority classification (High, Medium, Low) for restoration based on the relative contribution of foliar CH_4_ oxidation

| Species | Mean CH<sub>4</sub> oxidation<br>(nmol CO<sub>2</sub>-eq m<sup>-2</sup> s<sup>-1</sup>) | Mean N<sub>2</sub>O emission<br>(nmol CO<sub>2</sub>-eq m<sup>-2</sup> s<sup>-1</sup>) | CH<sub>4</sub>-to-N<sub>2</sub>O<br>impact ratio | Priority |
| --- | --- | --- | --- | --- |
| *Q. macrocarpa* | 24.8 | 0.0435 | 1135 | High |
| *A. saccharum* | 51.9 | 0.0995 | 592 | High |
| *U. americana* | 6.67 | 0.0191 | 547 | High |
| *Q. palustris* | 18.2 | 0.0744 | 512 | High |
| *T. americana* | 68.8 | 0.190 | 475 | High |
| *S. fragilis* | 29.7 | 0.0774 | 474 | High |
| *C. speciosa* | 56.0 | 0.137 | 458 | High |
| *A. freemanii* | 25.6 | 0.0813 | 417 | High |
| *V. opulus* | 11.9 | 0.0479 | 333 | Medium |
| *G. dioicus* | 24.2 | 0.121 | 222 | Medium |
| *B. papyrifera* | 24.9 | 0.127 | 204 | Medium |
| *A. rubrum* | 50.6 | 0.385 | 138 | Low |
| *C. sericea* | 13.2 | 0.117 | 129 | Low |
| *P. pungens* | 9.66 | 0.112 | 109 | Low |
| *A. balsamea* | 9.90 | 0.263 | 99.7 | Low |
| *A. saccharinum* | 12.9 | 0.141 | 98.1 | Low |
| *P. strobus* | 49.6 | 0.538 | 95.8 | Low |
| *T. occidentalis* | 72.8 | 0.834 | 93.8 | Low |
| *P. glauca* | 38.7 | 0.517 | 79.8 | Low |
| *R. typhina* | 22.2 | 0.285 | 78.6 | Low |
| *F. americana* | 28.2 | 0.409 | 67.7 | Low |
| *Q. rubra* | 37.6 | 0.716 | 54.0 | Low |
| *P. virginiana* | 1.04 | 0.0485 | 25.4 | Low |
| *G. triacanthos* | 2.97 | 0.423 | 7.31 | Low |
| *P. tremuloides* | 4.11 | 1.12 | 3.75 | Low |

## DISCUSSION

This study characterizes foliar CH_4_ and N_2_O fluxes among temperate tree species, revealing seasonal patterns and large species differences relevant for GHG budgets. Key findings are: (1) tree foliage acted as a net CH_4_ sink in all species, with oxidation rates higher in fall and in shade-tolerant species; (2) foliage consistently emitted N_2_O, with greater emissions in shade-intolerant species and no significant seasonality; and (3) flux variation was primarily at the species level, independent of phylogeny, and correlated with both soil fluxes and leaf functional traits. Metagenomic analyses of high- and low-flux hardwoods revealed an order-of-magnitude higher methanotroph abundance in the high-oxidation species as well as distinct microbial community compositions.

Until recently accurate measurements of foliar CH_4_ and N_2_O fluxes have been challenging due to instrument limitations. Our study is consistent with recent surveys indicating that tree foliage is generally a CH_4_ sink in upland forests (Sundqvist *et al*., 2012; Gorgolewski *et al*., 2023; Karim *et al*., 2024). Minimal uptake or net emissions occur under flooded conditions (Pangala *et al*., 2017; Gorgolewski *et al*., 2023), though foliar uptake has been detected in mangrove forests (Chang-yi *et al*., 1998; He *et al*., 2019). Net foliar CH_4_ emissions have also been found in some tropical pioneer species (Karim *et al*., 2024), and in *Pinus sylvestris* (Tenhovirta *et al*., 2024). For N_2_O, Karim et al., (2024) documented foliar emissions across 40 tropical species in situ, and Qin et al., (2024) reported emissions in 25 predominantly temperate species using excised branches.

### Controls on foliar CH_4_ oxidation

The consistent detection of foliar CH_4_ uptake across temperate tree species confirms that foliage serves as a measurable biological sink for atmospheric CH_4_, consistent with Gorgolewski et al., (2023), who reported comparable uptake rates in a near-natural temperate forest. Seasonal and functional-group differences were pronounced. The ∼33% higher CH_4_ oxidation rate in fall than in spring likely reflects two mechanisms: a longer growing period that allows endophytic methanotrophs to develop larger, more active populations (Shen & Fulthorpe, 2015), and autumn declines in stomatal conductance and transpiration (Bauerle *et al*., 2012), reducing soil-derived CH_4_ transport to the leaf (Moisan *et al*., 2024).

Systematic differences in foliar CH_4_ uptake also occurred with respect to tree shade tolerance. Shade-intolerant species, generally corresponding to early-successional pioneers, exhibited lower CH_4_ oxidation, whereas late-successional, shade-tolerant species showed higher uptake. This pattern corresponds to a longer leaf lifespan (Seiwa *et al*., 2006) which may facilitate microbial colonization. A comparable pattern was found in wet-season measurements in a tropical forest in Bangladesh, where mid-to late-successional tree species showed low foliar CH_4_ uptake while pioneer species showed net foliar CH_4_ emissions (Karim *et al*., 2024). These patterns are consistent in indicating that successional strategy strongly influences CH_4_-oxidizing capacity, and that late-seral species have a greater foliar CH_4_ uptake capacity.

Foliar CH_4_ uptake was further influenced by belowground processes and was related to leaf traits. Foliar and soil CH_4_ fluxes were correlated, consistent with xylem transport of dissolved CH_4_. Leaf physiological traits contributed secondary effects: CH_4_ oxidation increased slightly with transpiration, also consistent with a role of stomatal diffusion and internal gas transport (Karim *et al*., 2025). Structural traits such as specific leaf area had little influence, implying that internal biochemical and microbial factors dominate, while stem hydrology may mediate vertical soil–leaf coupling.

Collectively, these findings indicate that foliar CH_4_ flux is controlled by environmental conditions as well as plant-level physiological and microbial processes. Seasonal variation, successional stage, species traits, and soil–leaf linkages jointly determine the strength and variability of this CH_4_ sink. Foliar methanotrophy thus operates as part of a connected plant– soil continuum rather than an isolated leaf process. Most variation in CH_4_ oxidation occurred among species, suggesting future studies should emphasize broader taxonomic coverage over increased replication within species.

### Drivers of foliar nitrous oxide emissions

In contrast to CH_4_, foliage consistently emitted N_2_O, with little seasonal variation. This stability, together with the positive correlation between foliar and soil N_2_O fluxes, indicates contributions from both soil and leaf processes. The most plausible mechanism is upward transport of dissolved N_2_O via the xylem (Machacova *et al*., 2019). N_2_O produced in soils through nitrification and denitrification dissolves in soil water, is absorbed by roots, and moves with the transpiration stream to leaves, where it diffuses through stomata (Karim *et al*., 2025). The soil–leaf relationship supports this transport-based coupling, while the dependence with transpiration suggests water flux enhances gas movement but plateaus at high stomatal conductance.

The positive association between leaf nitrogen content and N_2_O emission indicates an endogenous foliar pathway as well. This likely reflects partial nitrate assimilation, in which incomplete reduction of nitrate (NO_3_^−^) to ammonium (NH_4_^+^) via nitrite (NO_2_^−^) produces N_2_O. This process occurs primarily in mitochondria under hypoxia, where nitrite is reduced to nitric oxide (NO) and then converted to N_2_O by cytochrome c oxidase (CcO). Although Smart & Bloom, (2001) suggested nitrite reductase involvement, later work indicates that N_2_O production occurs independently of this enzyme, even in nitrite reductase-deficient plants (Timilsina *et al*., 2020).

Shade-intolerant species had higher N_2_O emissions than shade-tolerant, likely reflecting greater foliar nitrogen assimilation. Soil microbial production provides a continuous N_2_O source transported via transpiration, while internal nitrate reduction contributes a parallel, plant-mediated source dependent on nitrogen status and metabolism. The lack of seasonal variation suggests both pathways are governed by relatively stable processes: sustained soil microbial activity under moderate moisture and temperature, and persistent internal nitrogen metabolism.

### Methanotrophic guilds and functional gene abundance in relation to foliar CH_4_ oxidation

We observed that the difference in CH_4_ uptake between *T. americana* and *P. virginiana* corresponded to distinct foliar methanotroph communities. In *T. americana*, high CH_4_ uptake was associated with Type I methanotrophs (*Methylococcales*), particularly *Candidatus Methylospira*, with *Methylomicrobium* and *Methylocaldum* at lower abundances. These taxa use the ribulose monophosphate (RuMP) pathway and encode particulate methane monooxygenase (pMMO) as their primary CH_4_-oxidizing enzyme (Trotsenko & Murrell, 2008). Although their CH_4_ affinity is low to moderate (AlSayed *et al*., 2018), high cell densities sustain measurable oxidation under CH_4_ limitation. The pmoA sequencing yielded 29,000–48,000 high-quality reads, over an order of magnitude higher than in *P. virginiana*, with dominant ASVs affiliated with uncultured *Methylococcales* carrying canonical pMMO genes. These results indicate that *T. americana* hosts a metabolically active Type I community with substantial CH_4_-oxidizing capacity.

In contrast, *P. virginiana* foliage had limited methanotrophic potential. Type I methanotrophs were nearly absent; facultative Type II methanotrophs (*Methylobacterium*– *Methylorubrum, Beijerinckiaceae*) were dominant. These organisms possess pmoA and oxidize C_1_ compounds but primarily rely on methylotrophy rather than CH_4_ oxidation (Haque *et al*., 2020). Rare *Methylocystis*-like sequences indicated sparse Type II methanotrophs with potential low- and high-affinity pMMO isozymes (Haque *et al*., 2020), but total pmoA reads were low, consistent with microbial limitation of net CH_4_ uptake. Facultative methanotrophs likely use methanol derived from plant processes, such as apoplastic pectin demethylation (Claudia *et al*., 2003; Dumont *et al*., 2014) as the primary substrate sustaining microbial growth.

Interpretation should consider molecular limitations and microbial interactions. The apparent absence of high-affinity methanotrophs (e.g., USCα) may reflect primer bias and database gaps (Claudia *et al*., 2003; Dumont *et al*., 2014). Methanotrophic communities in soils function as syntrophic consortia, where high-affinity methanotrophs oxidize CH_4_ to methanol, supporting facultative Type II populations (Zheng *et al*., 2024). Such cross-feeding may allow lower-affinity populations to persist under CH_4_-limited conditions and is likely relevant in foliar communities.

### Implications for global ecosystem budgets and future research

Temperate tree foliage functions as a dynamic biogeochemical interface that has not been fully integrated into global greenhouse gas budgets. Foliar CH_4_ uptake has been estimated to contribute ∼38% of daytime CH_4_ consumption in upland temperate forests (Gorgolewski *et al*., 2023). Our findings demonstrate that this foliar sink also exhibits marked seasonal variation, with oxidation increasing by ∼33% in the fall. Although foliar N_2_O emissions were observed in all species examined, preliminary GWP-weighted estimates indicate that CH_4_ uptake exerts the dominant climate influence (Table 4). Across all leaves, CH_4_ oxidation generates a climate impact approximately 250 times greater than foliar N_2_O emission, with functional type differences ranging from approximately 96 times in conifers to approximately 298 times in deciduous species (SM 12). These ratios reflect both the approximately 1000-fold difference in raw flux magnitude and the global warming potential of 27 for CH_4_ relative to 273 for N_2_O.

Previous efforts to upscale foliar CH_4_ and N_2_O fluxes to global budgets have been limited by narrow species representation and an absence of in situ measurements (Bloom *et al*., 2010; Qin *et al*., 2024), reducing confidence in global estimates. The present study addresses this gap by providing ecosystem-resolved flux measurements across multiple species while simultaneously identifying the mechanistic drivers of variation. While CH_4_ oxidation is consistently the dominant component of foliar greenhouse gas exchange, its magnitude varies substantially among species. Impact ratios span from 3.8 in *P. tremuloides* to 1135 in *Q. macrocarpa*, with *Q. macrocarpa* and *Acer saccharum* exerting disproportionately strong CH_4_ mitigation effects. This variation, spanning more than three orders of magnitude, demonstrates the critical role of species identity in shaping canopy-level greenhouse gas dynamics. Incorporating species-resolved flux estimates into Earth system models will improve projections of ecosystem greenhouse gas budgets and can potentially inform restoration planning that prioritizes high-oxidation species to enhance climate mitigation.

Our findings confirm that foliar CH_4_ oxidation is widespread among temperate tree species in upland North American forests, including within restoration plantings. Seasonal increases in oxidation during autumn and elevated methanotroph abundance in *T. americana* relative to very low abundance in *P. virginiana* suggest that microbial colonization limits the magnitude of foliar CH_4_ uptake. Although some earlier studies reported low CH_4_ oxidation or net CH_4_ efflux in conifers (Gorgolewski *et al*., 2023; Kohl *et al*., 2023; Tenhovirta *et al*., 2024), we observed higher mean oxidation rates in evergreen species compared to hardwoods, consistent with a colonization limitation mechanism. N_2_O emissions remained small relative to CH_4_ uptake across all species, reinforcing the dominant climate role of CH_4_ fluxes.

These patterns have clear implications for forest restoration and management. Species characterized by high foliar CH_4_ oxidation can be strategically prioritized to enhance greenhouse gas mitigation benefits. Key research needs include determining whether similar methanotrophic taxa are active across species, elucidating the pathways of microbial colonization, and assessing whether leaf-associated methanotrophs influence carbon and nitrogen cycling in litter and soil following leaf fall.

## ACKNOWLEDGEMENTS

This research was supported by the Natural Sciences and Engineering Research Council of Canada (NSERC). We thank Imrul Kayes, Liam Douglas, Jennifer Barrett, Jovana Shrestha, Melanie Sifton, Rachael Harman-Denhoed, Md Abdul Halim, and Rokaiya Binte Zahir for their technical assistance and contributions to field measurements, soil analyses, and site documentation. We also acknowledge Metagenom Bio Life Science Inc. (Waterloo, Canada) for DNA sequencing support and the Downsview Park authority (Ontario, Canada) for granting field access.

## AUTHOR CONTRIBUTIONS

M.R.K. conceptualized the study, collected and curated data, performed analyses, developed methodology and software. S.C.T. provided supervision, secured funding, offered conceptual guidance. M.R.K. wrote the first manuscript draft with contributions from S.C.T., who also reviewed and edited the manuscript.

## COMPETING INTERESTS

The authors declare no competing interests.

### DATA AVAILABILITY

The source data used to generate all graphs and charts presented in this study are publicly available via the Scholars Portal Dataverse repository at: https://doi.org/10.5683/SP3/LSD4IH

## REFERENCE

Ainsworth EA, Rogers A. 2007. The response of photosynthesis and stomatal conductance to rising [CO_2_]: mechanisms and environmental interactions. Plant, Cell & Environment 30: 258–270.

AlSayed A, Fergala A, Khattab S, Eldyasti A. 2018. Kinetics of type I methanotrophs mixed culture enriched from waste activated sludge. Biochemical Engineering Journal 132: 60–67.

Asner GP, Scurlock JMO, A. Hicke J. 2003. Global synthesis of leaf area index observations: implications for ecological and remote sensing studies. Global Ecology and Biogeography 12: 191–205.

Aurangojeb M, Klemedtsson L, Rütting T, He H, Weslien P, Banzhaf S, Kasimir A. 2017. Nitrous oxide emissions from Norway spruce forests on drained organic and mineral soil. Canadian Journal of Forest Research 47: 1482–1487.

Baldrian P. 2017. Forest microbiome: diversity, complexity and dynamics. FEMS Microbiology Reviews 41: 109–130.

Barba J, Bradford MA, Brewer PE, Bruhn D, Covey K, van Haren J, Megonigal JP, Mikkelsen TN, Pangala SR, Pihlatie M, et al. 2019. Methane emissions from tree stems: a new frontier in the global carbon cycle. New Phytologist 222: 18–28.

Bates D, Mächler M, Bolker B, Walker S. 2015. Fitting Linear Mixed-Effects Models Using lme4. Journal of Statistical Software 67: 1–48.

Bauerle WL, Oren R, Way DA, Qian SS, Stoy PC, Thornton PE, Bowden JD, Hoffman FM, Reynolds RF. 2012. Photoperiodic regulation of the seasonal pattern of photosynthetic capacity and the implications for carbon cycling. Proceedings of the National Academy of Sciences 109: 8612–8617.

Bloom AA, Lee-Taylor J, Madronich S, Messenger DJ, Palmer PI, Reay DS, McLeod AR. 2010. Global methane emission estimates from ultraviolet irradiation of terrestrial plant foliage. New Phytologist 187: 417–425.

Burghardt W, Morel JL, Zhang G-L. 2015. Development of the soil research about urban, industrial, traffic, mining and military areas (SUITMA). Soil Science and Plant Nutrition 61: 3–21.

Callahan BJ, McMurdie PJ, Rosen MJ, Han AW, Johnson AJA, Holmes SP. 2016. DADA2: High-resolution sample inference from Illumina amplicon data. Nature Methods 13: 581–583.

Chang-yi L, Yuk-shan W, Nora F. Y. T, Yong Y, Sheng-hui C, Peng L. 1998. Preliminary studies on methane fluxes in Hainan mangrove communities. Chinese Journal of Oceanology and Limnology 16: 64–71.

Claudia K, André L, Dunfield PF. 2003. Diversity and activity of methanotrophic bacteria in different upland soils. Applied and Environmental Microbiology 69: 6703–6714.

Covey KR, Megonigal JP. 2019. Methane production and emissions in trees and forests. New Phytologist 222: 35–51.

Crins WJ, Gray PA, Uhlig PWC, Wester MC. 2009. The ecosystems of Ontario, Part 1: ecozones and ecoregions. Ontario, CA: Ontario Ministry of Natural Resources.

De Sousa CA, Spiess TB. 2018. The management of brownfields in Ontario: A comprehensive review of remediation and reuse characteristics, trends, and outcomes, 2004–2015. Environmental Practice 20: 4–15.

Downsview Park. 2025. Wildlife at Downsview Park.

Dueck TA, De Visser R, Poorter H, Persijn S, Gorissen A, De Visser W, Schapendonk A, Verhagen J, Snel J, Harren FJM, et al. 2007. No evidence for substantial aerobic methane emission by terrestrial plants: a 13C-labelling approach. New Phytologist 175: 29–35.

Dumont MG, Lüke C, Deng Y, Frenzel P. 2014. Classification of pmoA amplicon pyrosequences using BLAST and the lowest common ancestor method in MEGAN. Frontiers in Microbiology 5: 34.

Engineer CB, Hashimoto-Sugimoto M, Negi J, Israelsson-Nordström M, Azoulay-Shemer T, Rappel WJ, Iba K, Schroeder JI. 2016. CO_2_ sensing and CO_2_ regulation of stomatal conductance: Advances and open questions. Trends in Plant Science 21: 16–30.

Environment Canada. 2025. Canadian Climate Normals 1991-2020. Environment and natural resources.

FAO. 2014. World reference base for soil resources 2014: International soil classification system for naming soils and creating legends for soil maps. Rome: FAO.

Galbraith JM. 2018. Human-altered and human-transported (HAHT) soils in the U.S. soil classification system. Soil Science and Plant Nutrition 64: 190–199.

Garnier E, Shipley B, Roumet C, Laurent G. 2001. A standardized protocol for the determination of specific leaf area and leaf dry matter content. Functional Ecology 15: 688–695.

Gastauer M, Meira-Neto JAA. 2016. An enhanced calibration of a recently released megatree for the analysis of phylogenetic diversity. Brazilian Journal of Biology 76: 619–628.

Gauci V, Pangala SR, Shenkin A, Barba J, Bastviken D, Figueiredo V, Gomez C, Enrich-Prast A, Sayer E, Stauffer T, et al. 2024. Global atmospheric methane uptake by upland tree woody surfaces. Nature 631: 796–800.

Genco T. 2007. Downsview park: city planning through the development of a model sustainable community. WIT Transactions on Ecology and the Environment 102: 209–218.

Gorgolewski AS, Caspersen JP, Vantellingen J, Thomas SC. 2023. Tree foliage is a methane sink in upland temperate forests. Ecosystems 26: 174–186.

Gorgolewski AS, Vantellingen J, Caspersen JP, Thomas SC. 2022. Overlooked sources of methane emissions from trees: branches and wounds. Canadian Journal of Forest Research 52: 1165–1175.

Halim MA, Bieser JMH, Thomas SC. 2024. Large, sustained soil CO_2_ efflux but rapid recovery of CH_4_ oxidation in post-harvest and post-fire stands in a mixedwood boreal forest. Science of The Total Environment 930: 172666.

Haque MFU, Xu H-J, Murrell JC, Crombie A. 2020. Facultative methanotrophs – diversity, genetics, molecular ecology and biotechnological potential: a mini-review. Microbiology 166: 894–908.

He Y, Guan W, Xue D, Liu L, Peng C, Liao B, Hu J, Zhu Q, Yang Y, Wang X, et al. 2019. Comparison of methane emissions among invasive and native mangrove species in Dongzhaigang, Hainan Island. Science of The Total Environment 697: 133945.

Hoffmann M, Schulz-Hanke M, Garcia Alba J, Jurisch N, Hagemann U, Sachs T, Sommer M, Augustin J. 2017. A simple calculation algorithm to separate high-resolution CH_4_ flux measurements into ebullition- and diffusion-derived components. Atmos. Meas. Tech. 10: 109–118.

Iguchi H, Sato I, Sakakibara M, Yurimoto H, Sakai Y. 2012. Distribution of Methanotrophs in the Phyllosphere. Bioscience, Biotechnology, and Biochemistry 76: 1580–1583.

Iguchi H, Sato I, Yurimoto H, Sakai Y. 2013. Stress resistance and C_1_ metabolism involved in plant colonization of a methanotroph Methylosinus sp. B4S. Archives of Microbiology 195: 717–726.

Jeffrey LC, Maher DT, Chiri E, Leung PM, Nauer PA, Arndt SK, Tait DR, Greening C, Johnston SG. 2021. Bark-dwelling methanotrophic bacteria decrease methane emissions from trees. Nature Communications 12: 2127.

Karim MR, Halim MA, Thomas SC. 2024. Foliar methane and nitrous oxide fluxes in tropical tree species. Science of the Total Environment 954.

Karim MR, Halim MA, Thomas SC. 2025. Foliar methane and nitrous oxide fluxes in Salix bebbiana respond to light and soil factors. Communications Earth & Environment 6: 493.

Keppler F, Hamilton JTG, Braß M, Röckmann T. 2006. Methane emissions from terrestrial plants under aerobic conditions. Nature 439: 187–191.

Kikuzawa K. 2003. Phenological and morphological adaptations to the light environment in two woody and two herbaceous plant species. Functional Ecology 17: 29–38.

Kirschbaum MUF, Bruhn D, Etheridge DM, Evans JR, Farquhar GD, Gifford RM, Paul KI, Winters AJ. 2006. A comment on the quantitative significance of aerobic methane release by plants. Functional Plant Biology 33: 521–530.

Kohl L, Tenhovirta SAM, Koskinen M, Putkinen A, Haikarainen I, Polvinen T, Galeotti L, Mammarella I, Siljanen HMP, Robson TM, et al. 2023. Radiation and temperature drive diurnal variation of aerobic methane emissions from Scots pine canopy. Proceedings of the National Academy of Sciences 120: e2308516120.

Legein M, Smets W, Vandenheuvel D, Eilers T, Muyshondt B, Prinsen E, Samson R, Lebeer S. 2020. Modes of action of microbial biocontrol in the phyllosphere. Frontiers in Microbiology 11: 1619.

LI-COR. 2015. Using the LI-8100A Soil Gas Flux System and the LI-8150 Multiplexer. Nebraska, U.S.: LI-COR Biosciences.

Lima AB, Muniz AW, Dumont MG. 2014. Activity and abundance of methane-oxidizing bacteria in secondary forest and manioc plantations of Amazonian Dark Earth and their adjacent soils. Frontiers in Microbiology 5: 550.

Machacova K, Borak L, Agyei T, Schindler T, Soosaar K, Mander Ü, Ah-Peng C. 2021. Trees as net sinks for methane (CH_4_) and nitrous oxide (N_2_O) in the lowland tropical rain forest on volcanic Réunion Island. New Phytologist 229: 1983–1994.

Machacova K, Vainio E, Urban O, Pihlatie M. 2019. Seasonal dynamics of stem N_2_O exchange follow the physiological activity of boreal trees. Nature Communications 10: 4989.

Martin M. 2011. Cutadapt removes adapter sequences from high-throughput sequencing reads. EMBnet.journal 17: 10–12.

Masson-Delmotte V, Zhai P, Pirani A, Connors SL, Péan C, Berger S, Caud N, Chen Y, Goldfarb L, Gomis MI, et al. (Eds). 2021. Climate Change 2021: The Physical Science Basis. Contribution of Working Group I to the Sixth Assessment Report of the Intergovernmental Panel on Climate Change. Cambridge, United Kingdom and New York, NY, USA: Cambridge University Press.

Mohanty SR, Mahawar H, Bajpai A, Dubey G, Parmar R, Atoliya N, Devi MH, Singh AB, Jain D, Patra A, et al. 2023. Methylotroph bacteria and cellular metabolite carotenoid alleviate ultraviolet radiation-driven abiotic stress in plants. Frontiers in Microbiology 13: 899268.

Moisan MA, Lajoie G, Constant P, Martineau C, Maire V. 2024. How tree traits modulate tree methane fluxes: A review. Science of The Total Environment 940: 173730.

Moisan MA, Maire V, Isabelle J, Philippo D, Martineau C. 2025. Tissue humidity and pH as important species traits regulating tree methane emissions in floodplain wetland forests. New Phytologist 248: 1713–1727.

Murali A, Bhargava A, Wright ES. 2018. IDTAXA: a novel approach for accurate taxonomic classification of microbiome sequences. Microbiome 6: 140.

Niinemets Ü, Valladares F. 2006. Tolerance to shade, drought, and waterlogging of temperate northern hemisphere trees and shrubs. Ecological Monographs 76: 521–547.

Nisbet EG, Dlugokencky EJ, Fisher RE, France JL, Lowry D, Manning MR, Michel SE, Warwick NJ. 2021. Atmospheric methane and nitrous oxide: challenges along the path to Net Zero. Philosophical Transactions of the Royal Society A: Mathematical, Physical and Engineering Sciences 379: 20200457.

Pachauri RK, Allen MR, Barros VR, Broome J, Cramer W, Christ R, Church JA, Clarke L, Dahe Q, Dasgupta P, et al. 2014. Climate Change 2014: Synthesis Report. Contribution of Working Groups I, II and III to the Fifth Assessment Report of the Intergovernmental Panel on Climate Change (RK Pachauri and L Meyer, Eds). Geneva, Switzerland: IPCC.

Pagel M. 1999. Inferring the historical patterns of biological evolution. Nature 401: 877–884.

Pangala SR, Enrich-Prast A, Basso LS, Peixoto RB, Bastviken D, Hornibrook ERC, Gatti LV, Marotta H, Calazans LSB, Sakuragui CM, et al. 2017. Large emissions from floodplain trees close the Amazon methane budget. Nature 552: 230–234.

Pinheiro J, Bates D, Team RC. 2024. Nlme: Linear and nonlinear mixed effects models.

Pouyat RV, Yesilonis ID, Russell-Anelli J, Neerchal NK. 2007. Soil chemical and physical properties that differentiate urban land-use and cover types. Soil Science Society of America Journal 71: 1010– 1019.

Prather MJ, Hsu J, DeLuca NM, Jackman CH, Oman LD, Douglass AR, Fleming EL, Strahan SE, Steenrod SD, Søvde OA, et al. 2015. Measuring and modeling the lifetime of nitrous oxide including its variability. Journal of Geophysical Research: Atmospheres 120: 5693–5705.

Putkinen A, Siljanen HMP, Laihonen A, Paasisalo I, Porkka K, Tiirola M, Haikarainen I, Tenhovirta S, Pihlatie M. 2021. New insight to the role of microbes in the methane exchange in trees: evidence from metagenomic sequencing. New Phytologist 231: 524–536.

Qin S, Pang Y, Hu H, Liu T, Yuan D, Clough T, Wrage-Mönnig N, Luo J, Zhou S, Ma L, et al. 2024. Foliar N_2_O emissions constitute a significant source to atmosphere. Global Change Biology 30: e17181.

Quast C, Pruesse E, Yilmaz P, Gerken J, Schweer T, Yarza P, Peplies J, Glöckner FO. 2013. The SILVA ribosomal RNA gene database project: improved data processing and web-based tools. Nucleic Acids Research 41: D590–D596.

R Core Team. 2024. R: A Language and Environment for Statistical Computing.

Revell LJ. 2024. phytools 2.0: an updated R ecosystem for phylogenetic comparative methods (and other things). PeerJ 12: e16505.

Santiago LS, Mulkey SS. 2003. A Test of Gas Exchange Measurements on Excised Canopy Branches of Ten Tropical Tree Species. Photosynthetica 41: 343–347.

Saurette D, Warren J, Heck R. 2021. Soils of Ontario. In: Krzic M, Walley FL, Diochon A, Paré MC, Farrell RE, eds. Digging into Canadian soils: An introduction to soil science. Pinawa, MB: Canadian Society of Soil Science.

Schoch CL, Ciufo S, Domrachev M, Hotton CL, Kannan S, Khovanskaya R, Leipe D, Mcveigh R, O’Neill K, Robbertse B, et al. 2020. NCBI Taxonomy: a comprehensive update on curation, resources and tools. Database 2020: baaa062.

Seiwa K, Kikuzawa K, Kadowaki T, Akasaka S, Ueno N. 2006. Shoot life span in relation to successional status in deciduous broad-leaved tree species in a temperate forest. New Phytologist 169: 537–548.

Shen SY, Fulthorpe R. 2015. Seasonal variation of bacterial endophytes in urban trees. Frontiers in Microbiology 6: 427.

Smart DR, Bloom AJ. 2001. Wheat leaves emit nitrous oxide during nitrate assimilation. Proceedings of the National Academy of Sciences 98: 7875–7878.

Sundqvist E, Crill P, Mölder M, Vestin P, Lindroth A. 2012. Atmospheric methane removal by boreal plants. Geophysical Research Letters 39: L21806.

Takahashi K, Kosugi Y, Kanazawa A, Sakabe A. 2012. Automated closed-chamber measurements of methane fluxes from intact leaves and trunk of Japanese cypress. Atmospheric Environment 51: 329– 332.

Tenhovirta SAM, Kohl L, Koskinen M, Polvinen T, Salmon Y, Paljakka T, Pihlatie M. 2024. Aerobic methane production in Scots pine shoots is independent of drought or photosynthesis. New Phytologist 242: 2440–2452.

Thomas SC, Bazzaz FA. 1999. Asymptotic height as a predictor of photosynthetic characteristics in Malaysian rain forest trees. Ecology 80: 1607–1622.

Timilsina A, Zhang C, Pandey B, Bizimana F, Dong W, Hu C. 2020. Potential pathway of nitrous oxide formation in plants. Frontiers in Plant Science 11: 1177.

Trotsenko YA, Murrell JC. 2008. Metabolic aspects of aerobic obligate methanotrophy. In: Advances in Applied Microbiology. Academic Press, 183–229.

Welch B, Gauci V, Sayer EJ. 2019. Tree stem bases are sources of CH_4_ and N_2_O in a tropical forest on upland soil during the dry to wet season transition. Global Change Biology 25: 361–372.

Welles JM, Demetriades-Shah TH, McDermitt DK. 2001. Considerations for measuring ground CO_2_ effluxes with chambers. Chemical Geology 177: 3–13.

Wikström N, Savolainen V, Chase MW. 2001. Evolution of the angiosperms: calibrating the family tree. Proceedings of the Royal Society of London. Series B: Biological Sciences 268: 2211–2220.

William W, Hyde Embriette R., Berg-Lyons Donna, Ackermann Gail, Humphrey Greg, Parada Alma, Gilbert Jack A., Jansson Janet K., Caporaso J. Gregory, Fuhrman Jed A., et al. 2015. Improved bacterial 16S rRNA gene (V4 and V4-5) and fungal internal transcribed spacer marker gene primers for microbial community surveys. mSystems 1: 10.1128/msystems.00009-15.

Yang M. 2022. Increases in the methane uptake of upland forest soil in China could significantly contribute to climate change mitigation. Forests 13: 1270.

Yu G, Smith DK, Zhu H, Guan Y, Lam TTY. 2017. ggtree: an r package for visualization and annotation of phylogenetic trees with their covariates and other associated data. Methods in Ecology and Evolution 8: 28–36.

Zhang L, Copini P, Weemstra M, Sterck F. 2016. Functional ratios among leaf, xylem and phloem areas in branches change with shade tolerance, but not with local light conditions, across temperate tree species. New Phytologist 209: 1566–1575.

Zheng Y, Cai Y, Jia Z. 2024. Role of methanotrophic communities in atmospheric methane oxidation in paddy soils. Frontiers in Microbiology 15: 1481044.

