## Supplementary material for "Species-level controls of foliar methane and nitrous oxide fluxes: roles of traits and microbes in temperate trees": Supplementary_downsview_np.docx

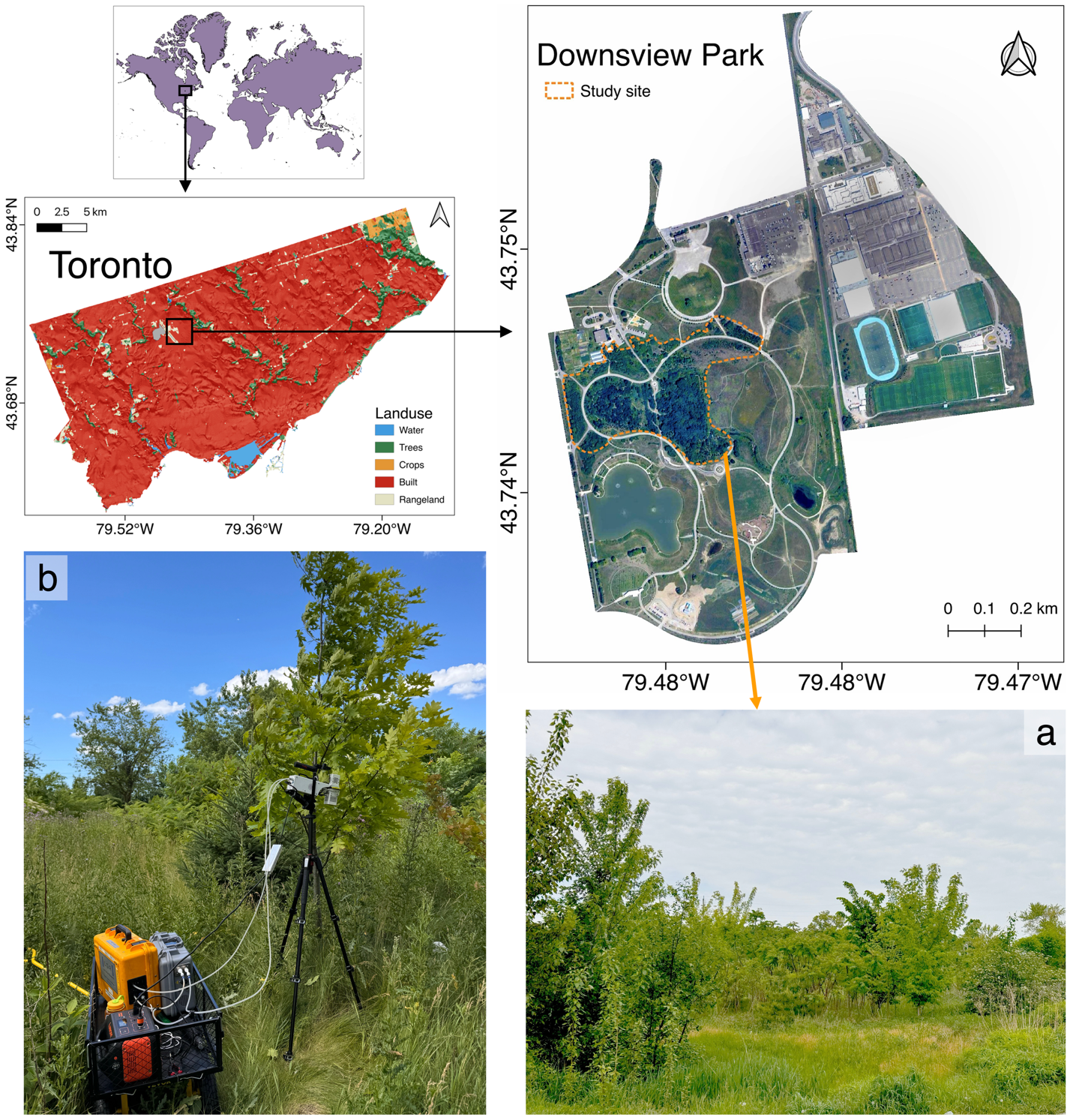

**SM 1:** Study site and greenhouse gas (GHG) flux measurement setup. The top panels show the study location in a global context (top left), land cover distribution in Toronto, Ontario, Canada (center left), and the specific boundaries of Downsview Park (right). Bottom panels illustrate (a) a general site view within Downsview Park and (b) the deployment of dynamic leaf chambers on trees for in-situ GHG flux measurements.

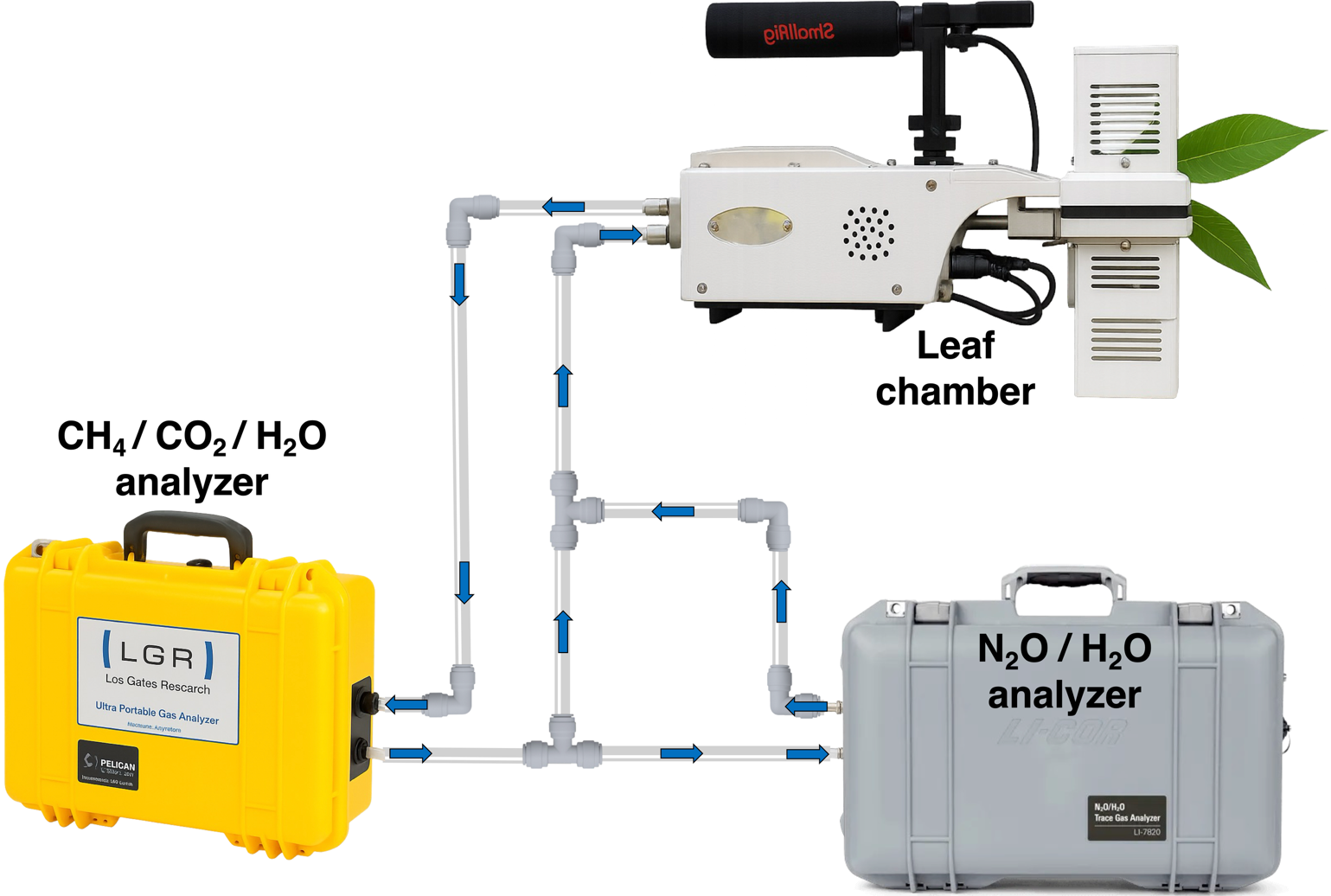

**SM 2:** Dynamic leaf chamber connected to trace gas analyzers for simultaneous in situ measurement of CH₄, N₂O, CO₂, and H₂O fluxes from intact foliage.

**SM 3**

**Sequencing quality and read processing summary**

Amplicon sequence quality and read retention were assessed using DADA2 v1.22 for both the 16S rRNA and pmoA datasets. The workflow included primer trimming, filtering, error correction, merging, and chimera removal. Read retention rates were calculated as the proportion of non-chimeric reads relative to the raw input.

Across all samples, 16S rRNA libraries showed consistently high data quality, with final non-chimeric read counts ranging from 31,242 to 246,502, corresponding to an average recovery of 92.3 ± 1.4%. The pmoA libraries exhibited greater variability, with 1,765–18,937 retained reads (20–73% recovery), reflecting expected differences in target gene abundance among species. Basswood (*Tilia americana*; samples 1–3) generally produced higher read depth and retention compared with Chokecherry (*Prunus virginiana*; samples 5–7), consistent with their distinct methanotrophic profiles.

**Table SM 3-1:** DADA2 read processing summary for 16S rRNA gene amplicons

| Sample | Raw (input) | Filtered | Denoised F | Denoised R | Merged | Non-chimeric | Retained (%) |
| --- | --- | --- | --- | --- | --- | --- | --- |
| Tilia-01 | 206,234 | 206,234 | 203,362 | 202,595 | 195,159 | 193,088 | 93.6 |
| Tilia-02 | 264,731 | 264,729 | 260,616 | 259,637 | 248,812 | 246,502 | 93.1 |
| Tilia-03 | 45,769 | 45,769 | 44,775 | 44,621 | 42,599 | 42,542 | 93.0 |
| Prunus-01 | 59,630 | 59,629 | 58,565 | 58,435 | 55,392 | 52,900 | 88.7 |
| Prunus-02 | 37,948 | 37,948 | 36,852 | 36,710 | 35,111 | 34,635 | 91.3 |
| Prunus-03 | 39,554 | 39,530 | 33,702 | 33,469 | 31,780 | 31,242 | 89.3 |

**Table SM 3-2:** DADA2 read processing summary for pmoA gene amplicons.

| Sample | Raw (reads.in) | Filtered | Denoised | Non-chimeric | Retained (%) |
| --- | --- | --- | --- | --- | --- |
| Tilia-01 | 25,927 | 25,686 | 19,189 | 18,937 | 73.0 |
| Tilia-02 | 8,398 | 8,226 | 3,467 | 3,467 | 41.3 |
| Tilia-03 | 8,263 | 7,996 | 1,780 | 1,765 | 21.4 |
| Prunus-01 | 13,672 | 13,149 | 116 | 116 | 0.9 |
| Prunus-02 | 12,495 | 12,090 | 66 | 66 | 0.5 |
| Prunus-03 | 10,550 | 10,167 | 95 | 95 | 0.9 |

In summary, the DADA2 pipeline achieved high-quality recovery for all 16S libraries and variable—but expected—retention for pmoA, confirming the robustness of sequence processing and supporting subsequent community composition analyses.

**SM 4:** Mean ± standard error (SE) of foliar CH₄ oxidation (nmol·m⁻²·s⁻¹) and N₂O emission (pmol·m⁻²·s⁻¹) across tolerance classes for shade, drought, and waterlogging. Values are shown separately for spring and fall 2024.

| Tolerance type | Response | Tolerance class | Spring 2024 | Fall 2024 |
| --- | --- | --- | --- | --- |
| **Shade tolerance** | **CH₄ oxidation (**nmol·m⁻²·s⁻¹**)** | 1 | 0.497 ± 0.102 | 0.754 ± 0.135 |
|  |  | 2 | 0.787 ± 0.081 | 1.10 ± 0.0914 |
|  |  | 3 | 0.945 ± 0.084 | 1.22 ± 0.0903 |
|  |  | 4 | 0.971 ± 0.103 | 1.34 ± 0.141 |
|  |  | 5 | 1.05 ± 0.193 | 1.24 ± 0.204 |
|  | **N₂O emission (p**mol·m⁻²·s⁻¹**)** | 1 | 2.15 ± 0.426 | 2.22 ± 0.467 |
|  |  | 2 | 0.863 ± 0.068 | 0.913 ± 0.078 |
|  |  | 3 | 0.982 ± 0.108 | 1.12 ± 0.116 |
|  |  | 4 | 0.742 ± 0.093 | 0.784 ± 0.088 |
|  |  | 5 | 0.559 ± 0.099 | 0.769 ± 0.122 |
| **Drought tolerance** | **CH₄ oxidation (**nmol·m⁻²·s⁻¹) | 1 | 0.590 ± 0.091 | 0.875 ± 0.116 |
|  |  | 2 | 0.846 ± 0.066 | 1.22 ± 0.074 |
|  |  | 3 | 1.02 ± 0.106 | 1.28 ± 0.124 |
|  |  | 4 | 1.06 ± 0.099 | 1.30 ± 0.098 |
|  |  | 5 | 0.104 ± 0.018 | 0.116 ± 0.025 |
|  | **N₂O emission (p**mol·m⁻²·s⁻¹) | 1 | 0.535 ± 0.101 | 0.713 ± 0.132 |
|  |  | 2 | 1.04 ± 0.115 | 1.15 ± 0.125 |
|  |  | 3 | 1.14 ± 0.127 | 1.22 ± 0.132 |
|  |  | 4 | 0.504 ± 0.062 | 0.570 ± 0.049 |
|  |  | 5 | 1.49 ± 0.147 | 1.61 ± 0.187 |
| **Waterlogging tolerance** | **CH₄ oxidation (**nmol·m⁻²·s⁻¹**)** | 1 | 1.26 ± 0.076 | 1.56 ± 0.085 |
|  |  | 2 | 0.342 ± 0.032 | 0.529 ± 0.043 |
|  |  | 3 | 0.668 ± 0.072 | 1.04 ± 0.094 |
|  |  | 4 | 0.882 ± 0.107 | 1.32 ± 0.085 |
|  | **N₂O emission (p**mol·m⁻²·s⁻¹) | 1 | 1.09 ± 0.088 | 1.19 ± 0.094 |
|  |  | 2 | 0.901 ± 0.186 | 1.06 ± 0.194 |
|  |  | 3 | 0.890 ± 0.076 | 0.958 ± 0.087 |
|  |  | 4 | 0.329 ± 0.048 | 0.238 ± 0.028 |

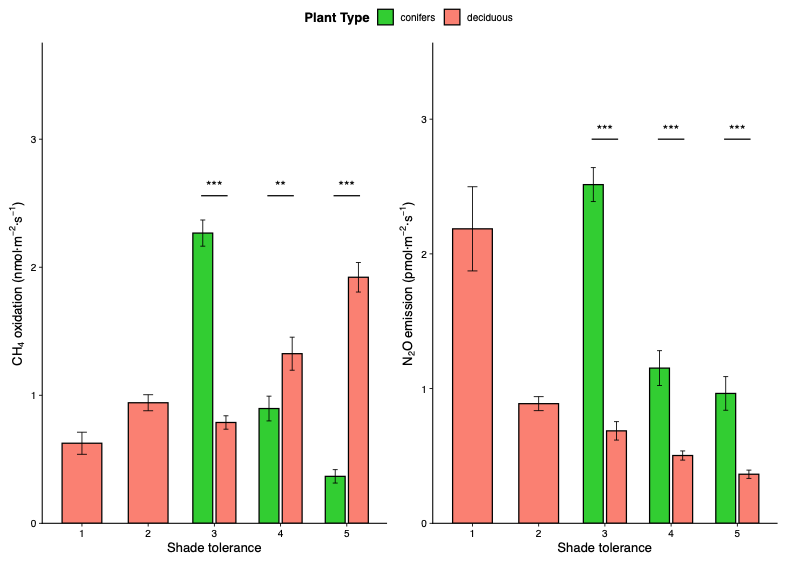

**SM 5**: Foliar CH₄ oxidation and N₂O emission across shade-tolerance classes in deciduous and coniferous species. Mean (± SE) foliar CH₄ oxidation (left) and N₂O emission (right) across shade-tolerance classes (1–5) for deciduous and coniferous trees. Bars represent species grouped by plant functional type, with significance indicators denoting pairwise differences between deciduous and coniferous species within each shade-tolerance class (t-tests; *p* < 0.05).

**SM 6**: Seasonal and species-type effects on foliar CO₂, CH₄, N₂O, and H₂O fluxes, based on linear mixed-effects models. ANOVA statistics (df, F-value, and p-value) indicate the significance of main and interactive effects between species type (deciduous vs. coniferous) and season (spring 2024 vs. fall 2024).

| Response variable | Fixed effect | Sum Sq | Mean Sq | Num  DF | DenDF | F value | P value |
| --- | --- | --- | --- | --- | --- | --- | --- |
| CH₄ oxidation (nmol·m⁻²·s⁻¹) | Type | 1.0766 | 1.0766 | 1 | 248.21 | 10.3175 | 0.0014** |
|  | Season | 4.7561 | 4.7561 | 1 | 248.77 | 45.5781 | <0.001*** |
|  | Type×Season | 0.4666 | 0.4666 | 1 | 248.77 | 4.4712 | 0.0354* |
| N₂O emission (pmol·m⁻²·s⁻¹) | Type | 2.21027 | 2.2102 | 1 | 248.21 | 26.1668 | <0.001*** |
|  | Season | 1.74155 | 1.7415 | 1 | 248.49 | 20.6178 | <0.001*** |
|  | Type×Season | 0.50712 | 0.5071 | 1 | 248.49 | 6.0037 | 0.0149* |
| H_2_O flux (mmol·m⁻²·s⁻¹) | Type | 3.6772 | 3.6772 | 1 | 248.07 | 6.1715 | 0.0136* |
|  | Season | 0.8729 | 0.8729 | 1 | 249.03 | 1.4650 | 0.2272 |
|  | Type×Season | 0.7114 | 0.7114 | 1 | 249.03 | 1.1940 | 0.2755 |
| CO_2_ uptake (µmol·m⁻²·s⁻¹) | Type | 154.926 | 154.926 | 1 | 248.29 | 32.9452 | <0.001*** |
|  | Season | 2.176 | 2.176 | 1 | 248.87 | 0.4627 | 0.4969 |
|  | Type×Season | 15.956 | 15.956 | 1 | 248.87 | 3.3931 | 0.0666 |

**SM 7:** Analysis of variance (ANOVA) results showing the effects of functional tolerance classes (drought and waterlogging) and season on foliar CH₄ oxidation and N₂O emission.

| Factor type | Response variable | Source of variation | Df | Sum Sq | Mean Sq | F value | Pr(>F) |
| --- | --- | --- | --- | --- | --- | --- | --- |
| **Drought tolerance** | CH₄ oxidation | Drought class | 4 | 24.49 | 6.123 | 9.817 | **<0.001** |
|  |  | Season | 1 | 10.80 | 10.798 | 17.312 | **<0.001** |
|  |  | Drought × Season | 4 | 0.78 | 0.196 | 0.314 | 0.868 |
|  | N₂O emission | Drought class | 4 | 35.8 | 8.961 | 8.402 | **<0.001** |
|  |  | Season | 1 | 1.2 | 1.211 | 1.136 | 0.287 |
|  |  | Drought × Season | 4 | 0.1 | 0.030 | 0.028 | 0.998 |
| **Waterlogging tolerance** | CH₄ oxidation | Waterlogging class | 3 | 81.09 | 27.030 | 53.374 | **<0.001** |
|  |  | Season | 1 | 10.80 | 10.798 | 21.322 | **<0.001** |
|  |  | Waterlogging × Season | 3 | 0.64 | 0.213 | 0.421 | 0.738 |
|  | N₂O emission | Waterlogging class | 3 | 15.4 | 5.134 | 4.653 | **0.003** |
|  |  | Season | 1 | 1.2 | 1.211 | 1.098 | 0.295 |
|  |  | Waterlogging × Season | 3 | 0.3 | 0.108 | 0.098 | 0.961 |

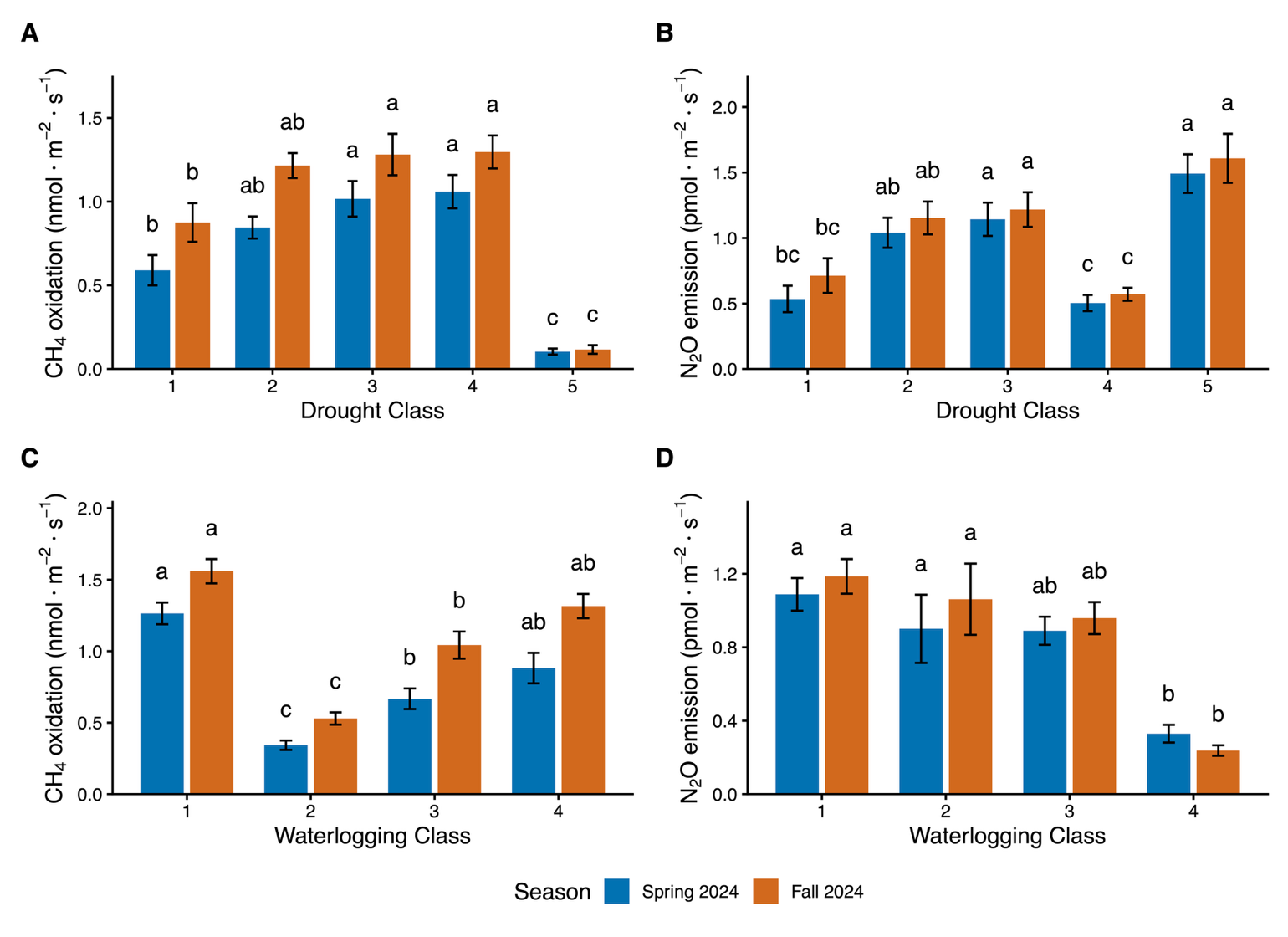

**SM 8:** Effects of functional tolerance classes (drought and waterlogging) on foliar CH₄ oxidation and N₂O emission across spring and fall seasons. Bars represent mean fluxes (± SE) for each tolerance class, with letters indicating significant differences based on Tukey’s HSD tests (*p* < 0.05). Increasing values along tolerance classes represent transitions from less to more stress-tolerant species.

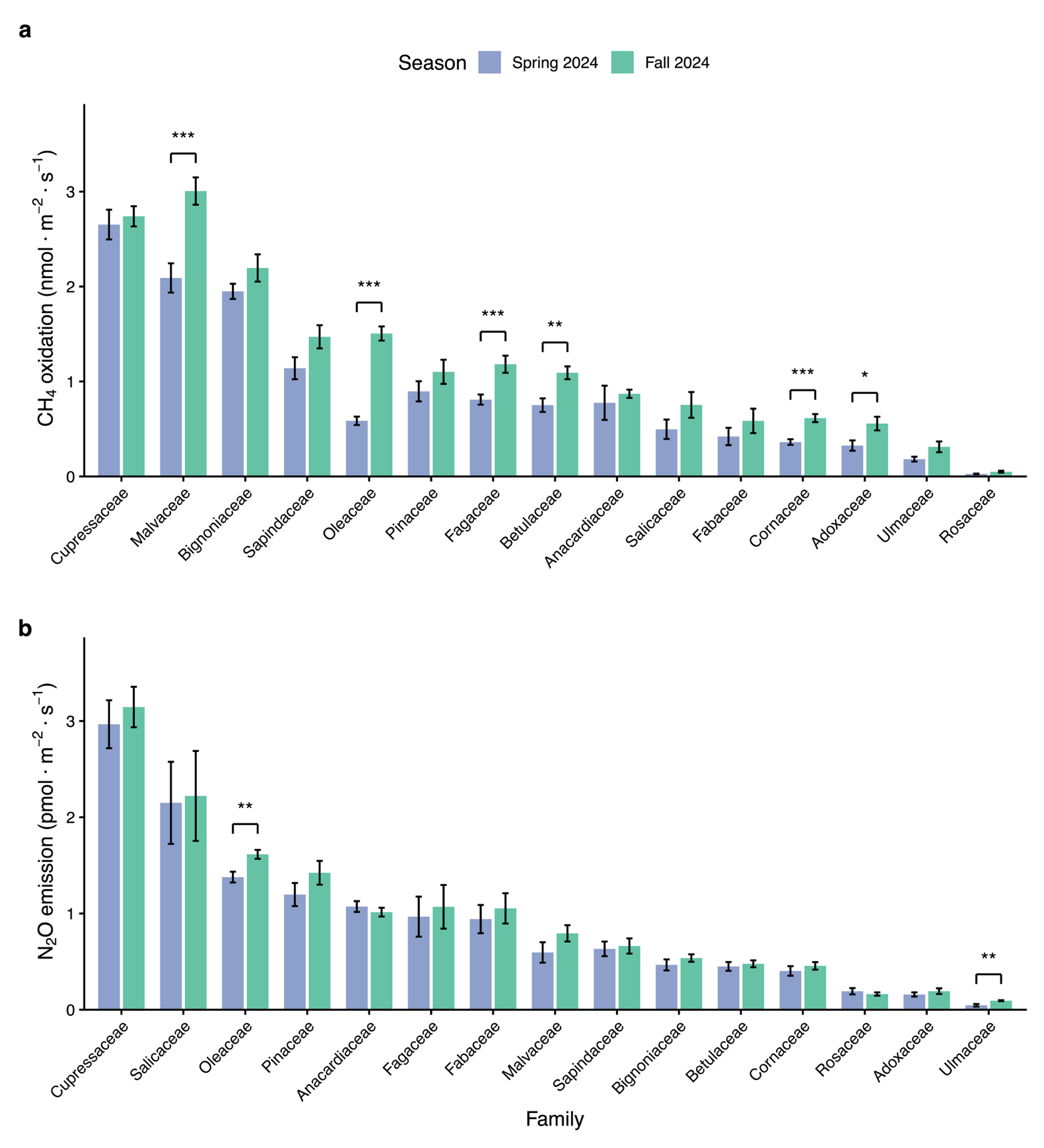

**SM 9:** Seasonal variation in foliar greenhouse gas fluxes across tree families. (a) Mean (± SE) rates of CH₄ oxidation (nmol·m⁻²·s⁻¹) and (b) mean (± SE) rates of N₂O emission (pmol·m⁻²·s⁻¹) measured on leaves during spring (blue) and fall (green) 2024 across major tree families. Families are ordered alphabetically; asterisks denote significant seasonal differences (**p* < 0.05, ***p* < 0.01, ****p* < 0.001).

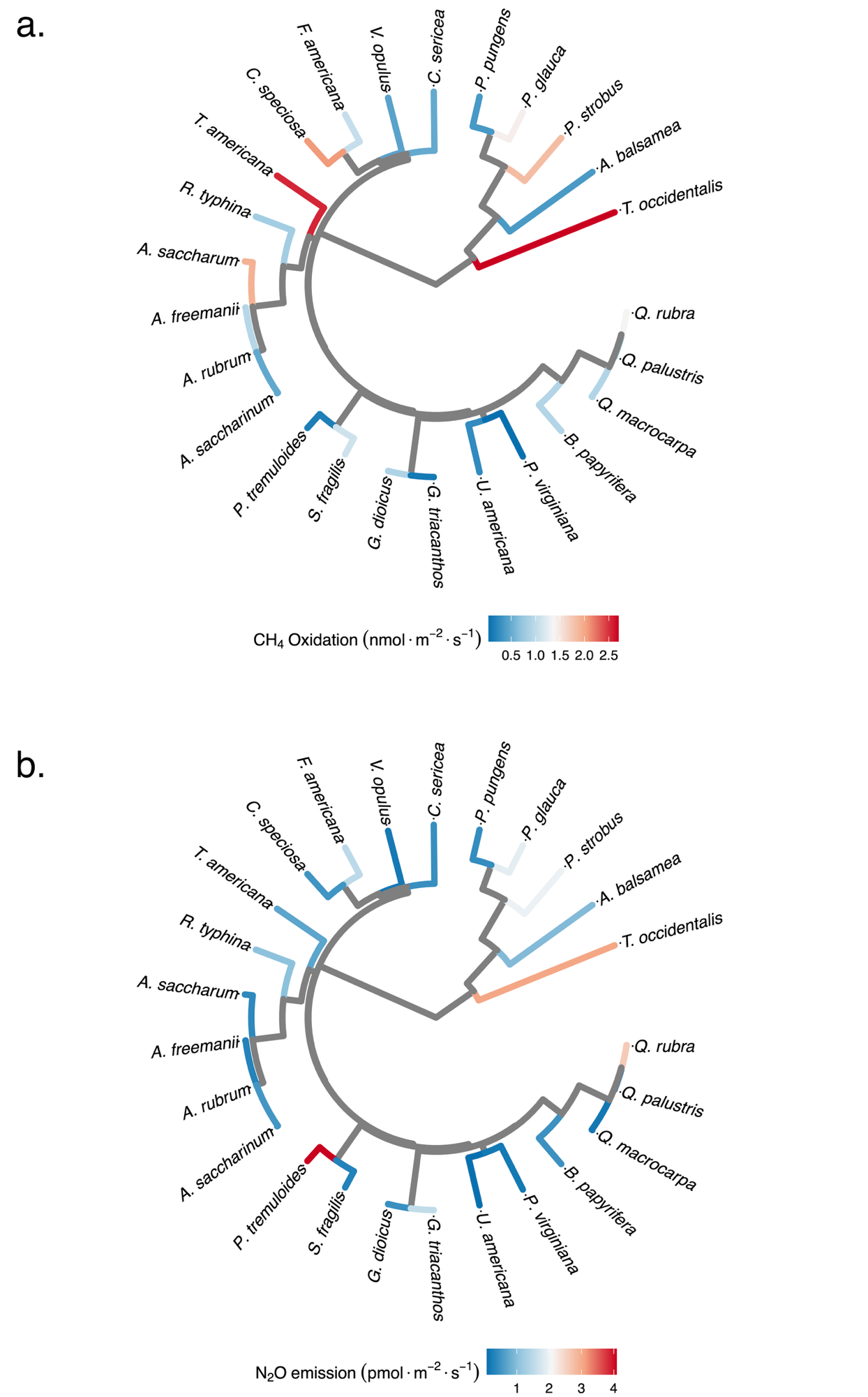

**SM 10:** Phylogenetic variation in foliar gas fluxes across temperate tree species. (a) Leaf CH₄ oxidation and (b) leaf N₂O emission. Tip colors on the phylogenetic tree represent the magnitude of each flux, mapped continuously along a gradient.

**SM 11:** Variance components, standard deviations (SD), and percentage of total variance explained for foliar CH₄ oxidation and N₂O fluxes at each hierarchical level.

| Leaf flux | Hierarchical level | Variance | SD | Variance explained (%) |
| --- | --- | --- | --- | --- |
| **CH₄** | Family | 8.70 × 10⁻¹⁰ | 2.95 × 10⁻⁵ | ≈0.00 |
|  | Species | 0.550 | 0.742 | 80.0 |
|  | Tree (individual) | 0.011 | 0.105 | 1.6 |
|  | Leaf | 2.68 × 10⁻¹⁰ | 1.64 × 10⁻⁵ | ≈0.00 |
|  | Residual | 0.124 | 0.352 | 18.0 |
| **N₂O** | Family | 2.69 × 10⁻⁸ | 1.64 × 10⁻⁴ | ≈0.00 |
|  | Species | 1.028 | 1.014 | 91.0 |
|  | Tree (individual) | 5.57 × 10⁻³ | 0.075 | 0.49 |
|  | Leaf | 6.02 × 10⁻³ | 0.078 | 0.53 |
|  | Residual | 0.092 | 0.303 | 8.1 |

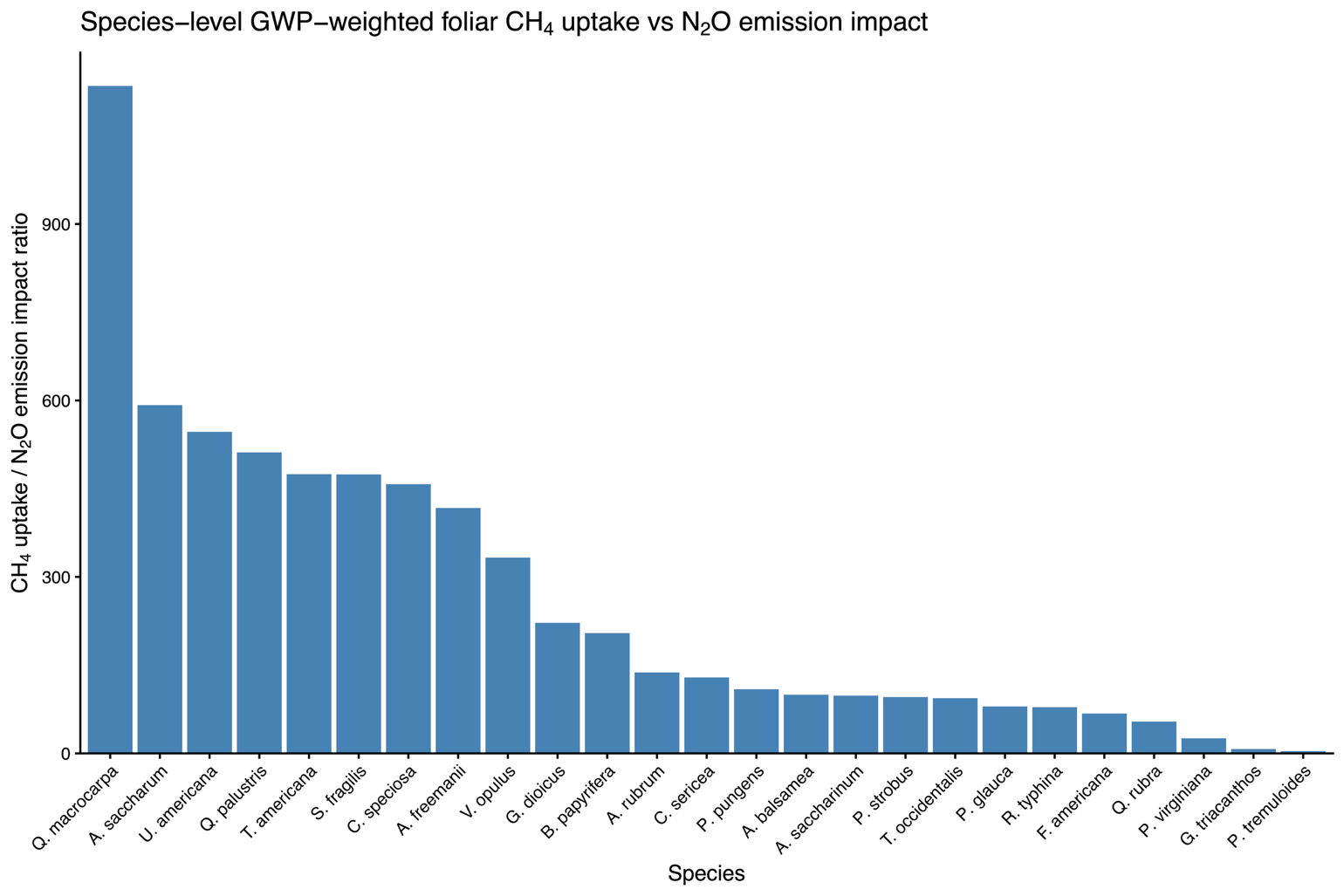

**SM 12.** Species-level comparison of the relative climate impact of foliar CH₄ uptake and N₂O emission. Bars represent the mean CH₄-to-N₂O impact ratio (GWP-weighted fluxes) for each species, ordered from highest to lowest. Values >1 indicate that foliar CH₄ oxidation dominates the net climate forcing compared with N₂O emission.
